# Tumour Extracellular Vesicles Induce Neutrophil Extracellular Traps To Promote Lymph Node Metastasis

**DOI:** 10.1101/2023.03.13.532413

**Authors:** Xin Su, Ariane Brassard, Alexandra Bartolomucci, Iqraa Dhoparee-Doomah, Qian Qiu, Thupten Tsering, Ramin Rohanizadeh, Olivia Koufos, Betty Giannias, France Bourdeau, Lixuan Feng, Sabrina Leo, Veena Sangwan, James Tankel, Daniela Quail, Jonathan Spicer, Julia V Burnier, Swneke Donovan Bailey, Lorenzo Ferri, Jonathan Cools-Lartigue

## Abstract

Lymph nodes (LNs) are frequently the first sites of metastasis. Currently, the only prognostic LN assessment is determining metastasis status. However, there is evidence suggesting that LN metastasis is facilitated by a pre-metastatic niche induced by tumour derived extracellular vehicles (EVs). Therefore, it is important to detect and modify the LN environmental changes. We have previously reported that neutrophil extracellular traps (NETs) can sequester and promote distant metastasis. Here, we first confirmed that LN NETs are associated with reduced patient survival. Next, we demonstrated that NETs deposition precedes LN metastasis and NETs inhibition abolishes LN metastases in animal mode. Furthermore, we discovered that EVs are essential to the formation of LN NETs. Lymphatic endothelial cells secrete CXCL8/2 in response to EVs inducing NETs formation and the promotion of LN metastasis. Our findings are the first to reveal the role of EV induced NETs in LN metastasis and provide potential immunotherapeutic vulnerabilities.

**Graphic Abstract:** Illustrative demonstration of the LNs premetastatic niche formation induced by EVs and NETs. Primary tumour constantly secretes EVs, which were actively uptaken by LECs. LECs subsequently secretes CXCL8 or CXCL2 upon EV reception. CXCL8 and CXCL2 are both neutrophil chemoattractants and potent NETs inducers. The following neutrophil recruitment and NETs formation lead to increased LN metastasis burden.

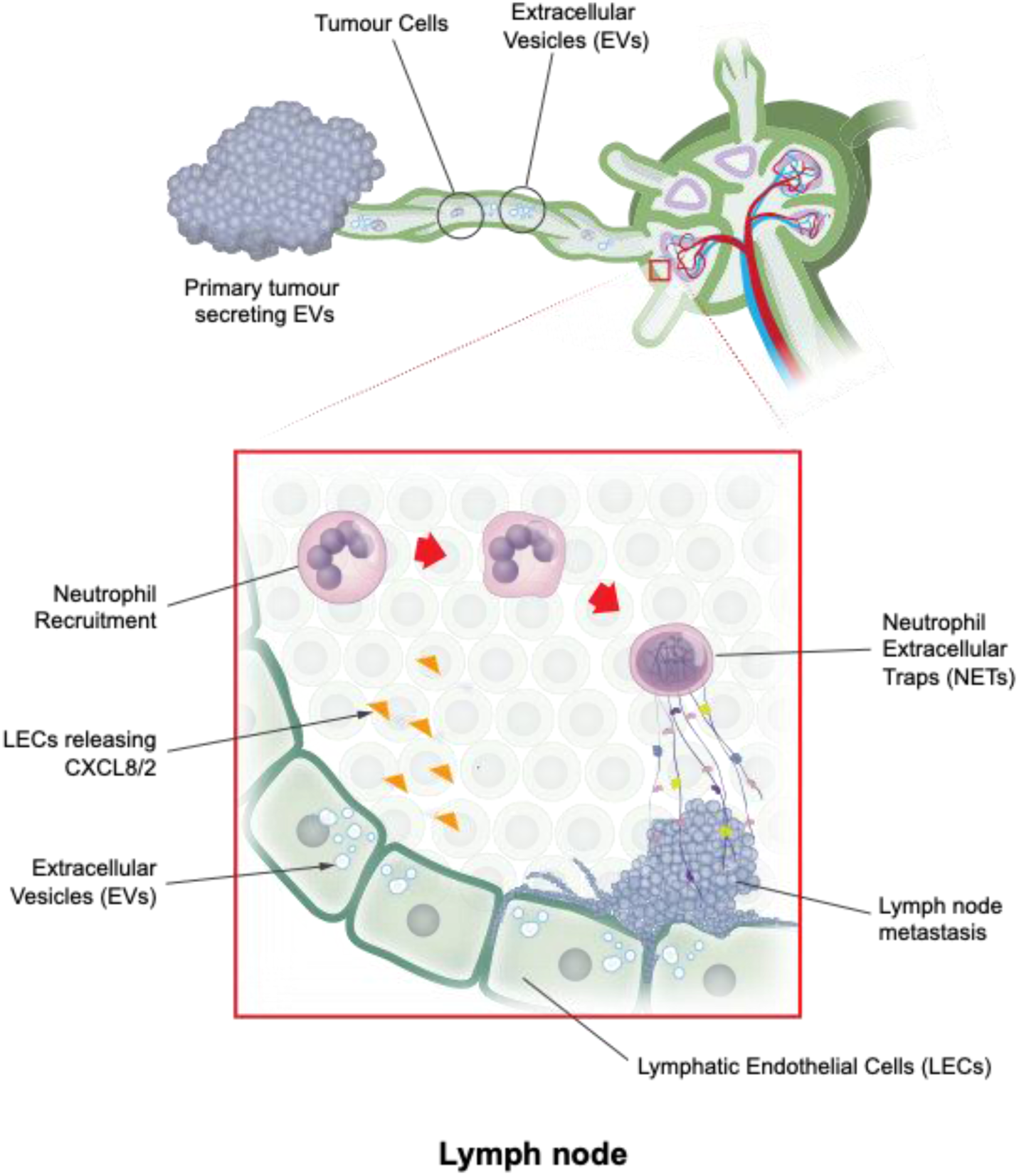

## Main

Solid organ tumours are treated with two goals in mind. The first is local tumour control for the purpose of symptom alleviation. The second is to prevent the development of distant metastasis, the major limitation to survival^1, 2^. Accordingly, whenever possible, primary tumours are excised along with regional lymph nodes (LN)^3, 4^. Regional node resection improves survival and allows for accurate disease staging^5^. The is of critical importance, as systemic therapies (e.g., cytotoxic chemotherapy, immunotherapy), are prescribed on the basis of risk factors for the development of fatal distant disease^6^. The presence of lymph node (LN) metastases is one of the strongest predictors of the subsequent development of fatal systemic disease^7^ and understanding the mechanisms is of major therapeutic interest^8^. Currently, the only relevant prognostic feature is the presence or absence of overt metastatic tumour cells within the LNs. However, this solely tumour centric system overlooks the role of immune and non-tumour compartment within the LNs^9^. This is critical, as no treatment strategy to date has reliably prevented the emergence of distant metastasis.

For metastatic colonisation to occur, the conditions within the LNs are optimised for tumour cell deposition and growth, a process known as pre-metastatic niche formation^10, 11^, which includes lymphoangiogenesis and the activation of various immune cells^12–14^. Neutrophils are resident cells within LNs^15^. Under normal physiologic condition, LN neutrophils play a role in host defence and present antigens^16–18^. However, in the context of malignancy, inflammation, defined as elevated neutrophil activity, and infiltration, are associated with reduced patient survival^19–21^. Inflammation is recognized as one of the hallmarks of cancer, facilitating functions necessary for tumour progression^22–24^. These include proliferation, resistance to therapy, and sequestration of tumour cells within distant organ sites^25–29^. Neutrophils are the most abundant leukocyte in humans and play a primary role in host defence to infection, however, in the context of malignancy, neutrophils are a major driver of inflammation^21, 30–32^. They mediate functions necessary for cancer progression, such as tumour cell proliferation, invasion, and metastasis^33–35^. Specifically, neutrophils can induce resistance to cytotoxic chemotherapy, radiotherapy, and immunotherapy^36–40^. Neutrophils can trap and support malignant cells within distant organs. This has been attributed to the elaboration of neutrophil extracellular traps (NETs)—extracellular strands of DNA decorated with biologically active peptides^16, 41^. These NETs play a role in host defence to infection and represent a primary effector function of neutrophils under normal physiologic conditions^42, 43^. Our group has played a leading role in demonstrating the role that NETs have in cancer progression and the profound abrogation of metastasis that occurs following NETs inhibition^37, 44, 45^. Since then, many of the mechanisms by which neutrophils facilitate tumour progression have been attributed to tumour-NETs interactions^46–48^.

Neutrophils exhibit great plasticity to environmental cues and the process of NETs release is believed to be mediated by tumour secretion of numerous kinds of factors, including extracellular vesicles (EVs)^36, 49, 50^. Tumour-derived EVs have been shown to be actively uptaken and transported by lymphatic vessels and are able to prepare a pre-metastatic niche in sentinel LNs for impending melanoma metastasis^51–53^. Moreover, EVs can polarize neutrophils from an anti-tumour (N1) phenotype to a pro-tumour (N2) phenotype^54^. EVs can also directly induce the formation of NETs^55^. However, it remains entirely unknown whether EV-mediated NETs production has a role in LN metastasis development. A better understanding of the microenvironmental changes initiating LN metastasis is needed to improve clinical understanding and intervention. Currently, no therapeutic modalities exist that selectively mitigate the consequences of neutrophil mediated inflammation.

In the current study, we demonstrate that lymphatic neutrophil accumulation and NETs deposition is associated with reduced survival in human gastroesophageal adenocarcinoma (GEA) patients. Furthermore, we demonstrate that lymphatic neutrophil accumulation and NETs deposition precedes and is required for LN metastatic outgrowth. Finally, we describe the role tumour derived EVs play in establishing a regional inflammatory microenvironment through the induction of lymphatic endothelial elaboration of CXCL8.

## Results

### Lymphatic NETs are associated with reduced survival in gastroesophageal cancer patients

Our group has previously demonstrated that neutrophil accumulation and NETs deposition promotes the development of metastasis at distant organ sites^37, 41, 44^. Here, we sought to determine if this process was conserved in LN metastasis, an earlier yet treatable stage during cancer progression. To characterise the pattern of lymphatic neutrophil recruitment and NETs deposition, we constructed tissue microarrays (TMAs) from 175 GEA surgical samples (Fig. 1a) (Table S1). The TMAs contained both patients with LN metastasis (N+) and without (N0). Within an N+ patient, not all nodes harbour metastatic cancer. Those that do are designated N+met, while those that do not are designated N+neg. Conversely, within an N0 patient, all regional nodes are free of cancer.

**Figure 1:**
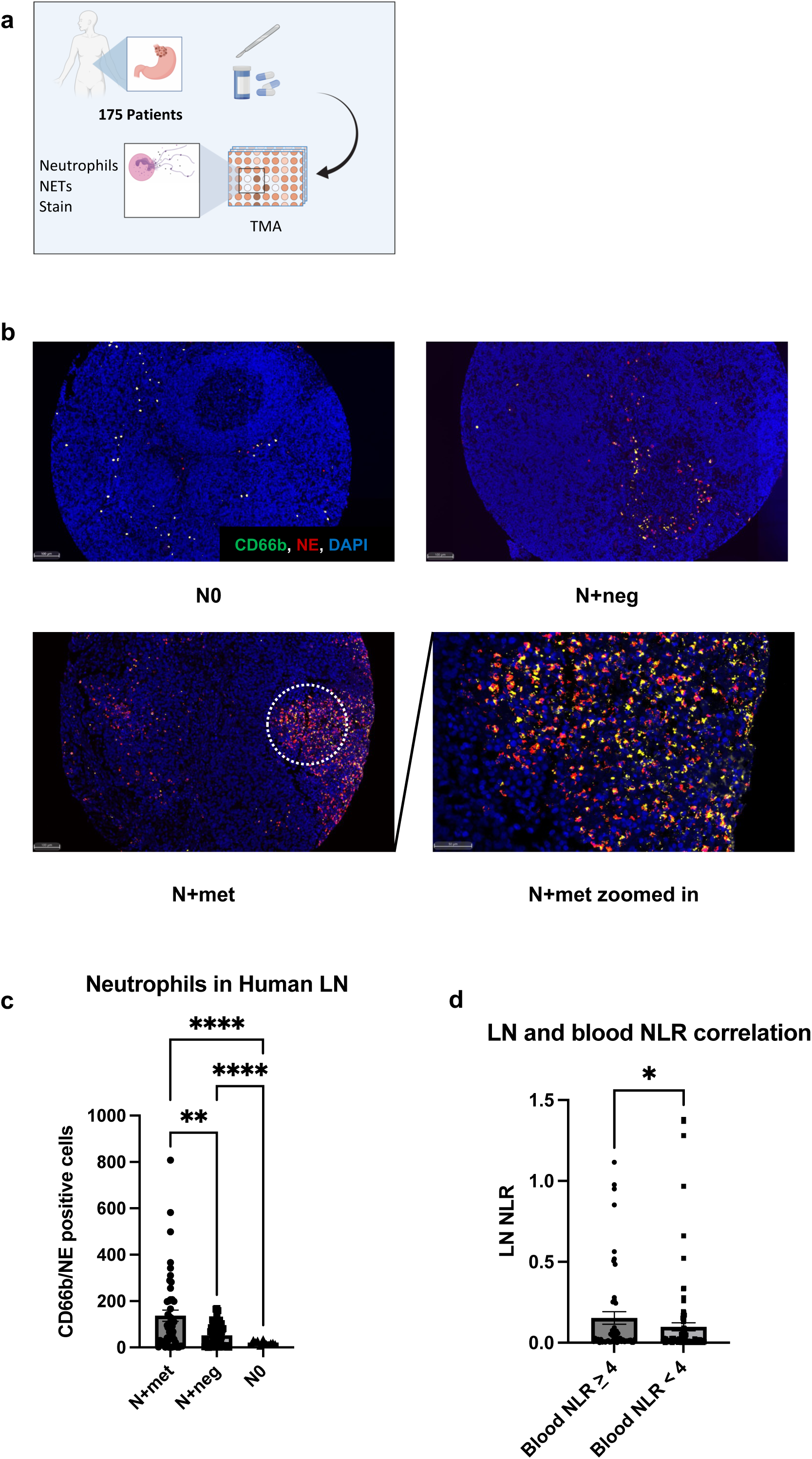
Neutrophils are recruited to regional lymph nodes (LNs) in gastroesophageal adenocarcinoma patients. (a) Schematic illustration of the TMA constructions (b) Representative images of lymphatic involvement by neutrophils depicting a node negative patient (N0, top left), tumour negative LN in a node positive patient (N+neg, top right), and a tumour positive LN (N+met, bottom left). Scale bars represent 100μm. White circle indicates the zoomed area shown in bottom right. Scale bars represent 50μm. (c) Quantification of lymphatic neutrophils (CD66b/NE positive cells) in N+met, N+neg and N0 LNs. (d) Comparison of LN neutrophil-lymphocyte ratio (LN NLR) with patient with high blood NLR (≥4) and patients with low blood NLR (<4). Data shown as mean ± SEM. n = 175; **, P < 0.01; **** P < 0.001 by Brown-Forsythe ANOVA test.

The TMAs stained with neutrophil markers CD66b and neutrophil elastase (NE), NETs marker citrullinated histone H3 (H3Cit) and GEA epithelial marker cytokeratin 7 (CK7). We found that neutrophils were present and progressively recruited to regional LNs in N+ patients (Fig. 1b). By quantification, we demonstrated that cancer positive nodes (N+met) have significantly higher neutrophil counts compared to cancer negative nodes (N+met vs. N+neg, 137.0 vs. 51.77, p = 0.0033) (N+met vs. N0, 137.0 vs. 21.23, p<0.0001) (Fig.1c). Notably, the neutrophil count in N+neg is significantly higher than within N0 (51.77 vs. 21.23, p<0.0001). This indicates that LN neutrophil recruitment may occur prior to lymphatic neoplastic invasion.

In addition, we assessed the relationship between the peripheral blood neutrophil-lymphocyte ratio (NLR), a standardized neutrophil quantification^19, 56^, and LN NLR (Fig. 1d). We found that patients with a blood NLR>4, which has previously been defined as elevated, also have a significantly higher LN NLR (0.02430 vs. 0.01374, p= 0.0451). This result suggests the dynamic correlation between lymphatic neutrophil accumulation and systemic inflammation.

Having shown that neutrophil accumulation within the regional LNs can occur alongside tumour LN infiltration, we therefore sought to determine if NETs deposition also takes place within the LN. Mirroring the same trend shown in nodal neutrophil infiltration, both N+met and N+neg nodes have higher levels of NETs deposition compared to N0 nodes (Fig. 2a, b). (0.05134 versus 0.1663, p = 0.0372) (0.05134 versus 0.1330, p = 0.0401) Moreover, we found that increased LN NETs quantity was significantly associated with reduced overall survival, both in N+met (hazard ratio [HR], 2.633, 604 versus 1415.5 days, p = 0.0014) and N+neg (hazard ratio [HR], 1.680, 688 versus 1161 days, p = 0.03) LNs (Fig. 2d).

**Figure 2:**
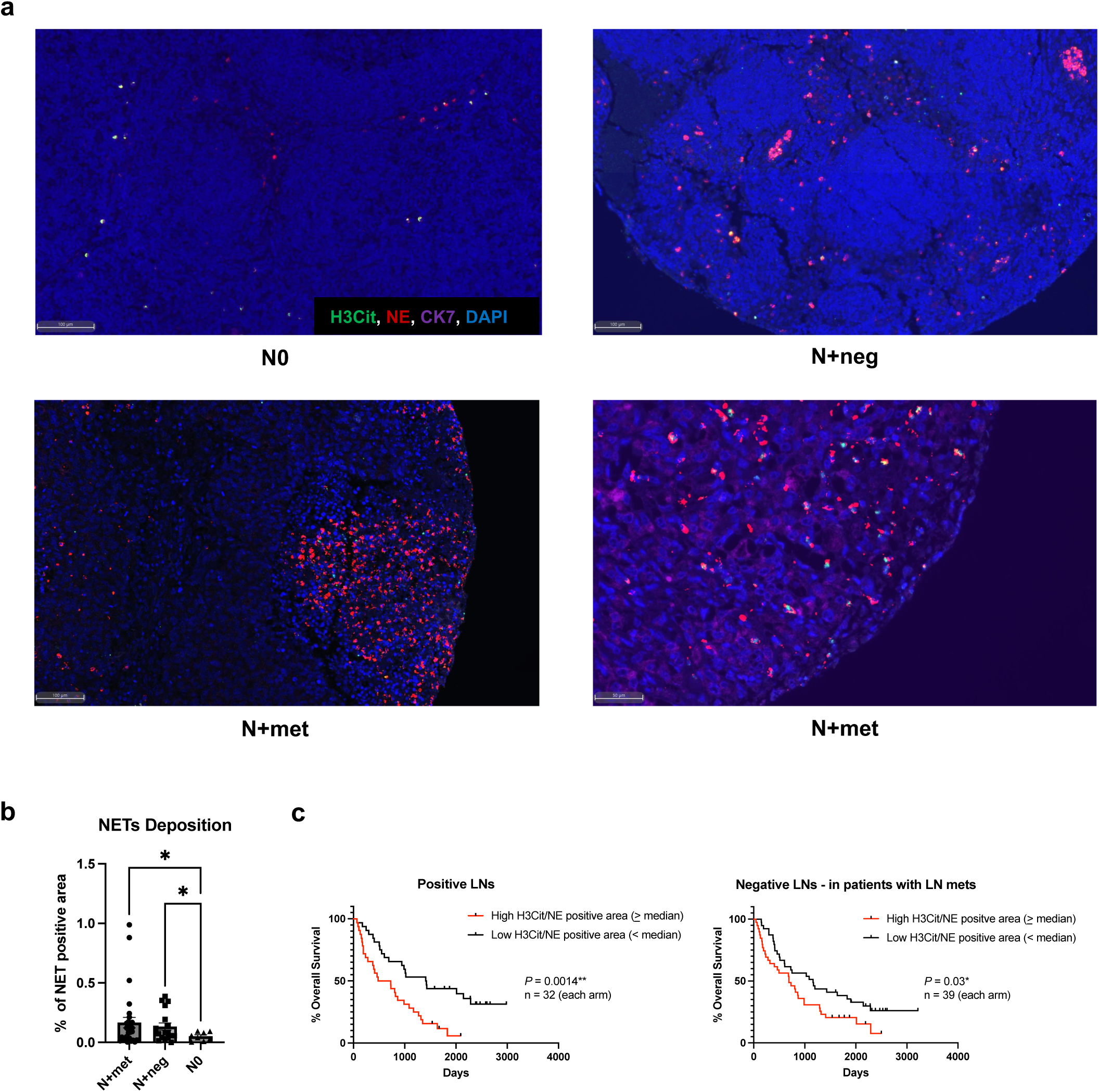
Elevated lymphatic NETs deposition is associated with poor survival. (a) Representative images of lymphatic NETs deposition pattern in N0 (top left), N+neg (top right), and N+met (bottom left) LNs. Scale bars represent 100μm. Zoomed details of NETs are shown in bottom right. Scale bars represent 50μm. (b)Quantification of lymphatic NETs (% of NETs positive area) in N+met, N+neg and N0 LNs. Data shown as mean ± SEM. n = 175; *, P < 0.05 by Brown-Forsythe ANOVA test. (c) Kaplan-Meier survival curves comparing survival of gastroesophageal patients with low versus high levels of median lymphatic NETs positive area, in both tumour positive and negative LNs. P value by Log-rank (Mantel-Cox) test.

Collectively, these results suggest a correlation between lymphatic neutrophil accumulation, NETs deposition, and ultimately survival in patients with GEA, which points to a pro-tumorigenic role of neutrophils and NETs in the progression of regional lymphatic disease. In addition, our data suggests that changes favourable for lymphatic metastasis may occur prior to nodal neoplastic ingress.

### Neutrophil accumulation and NETs deposition precedes overt lymphatic metastasis in vivo

The findings from GEA patients’ samples suggests that the changes favourable to lymphatic metastasis prior to nodal neoplastic ingress. Therefore, we developed an animal model in order to investigate the kinetics of LN NETs deposition. Briefly, two different cell lines, H59 and B16 F16 were injected into flank of C57bl/6 mice. Mice were sacrificed at day 7/10 and day 14 in order to reflect both a pre-metastatic and post-metastatic settings. We specifically used two different tumour cell lines to imply a universal biology and not cell specific phenomenology. Subsequently, the tumour draining ipsilateral inguinal LNs were resected and analysed (Fig.3a).

Using flow cytometry, we observed a significant increase in the percentage of viable neutrophils (Viability Dye eFlour780^-^CD11b^+^Ly6G^+^) in the premetastatic LNs of all tumour bearing mice (Fig. 3b). At this stage all LNs were devoid of neoplastic cells. At the post-metastatic stage, we observed a relative drop or stability of viable neutrophil numbers, for H59 and B16F10 cells, respectively.

**Figure 3:**
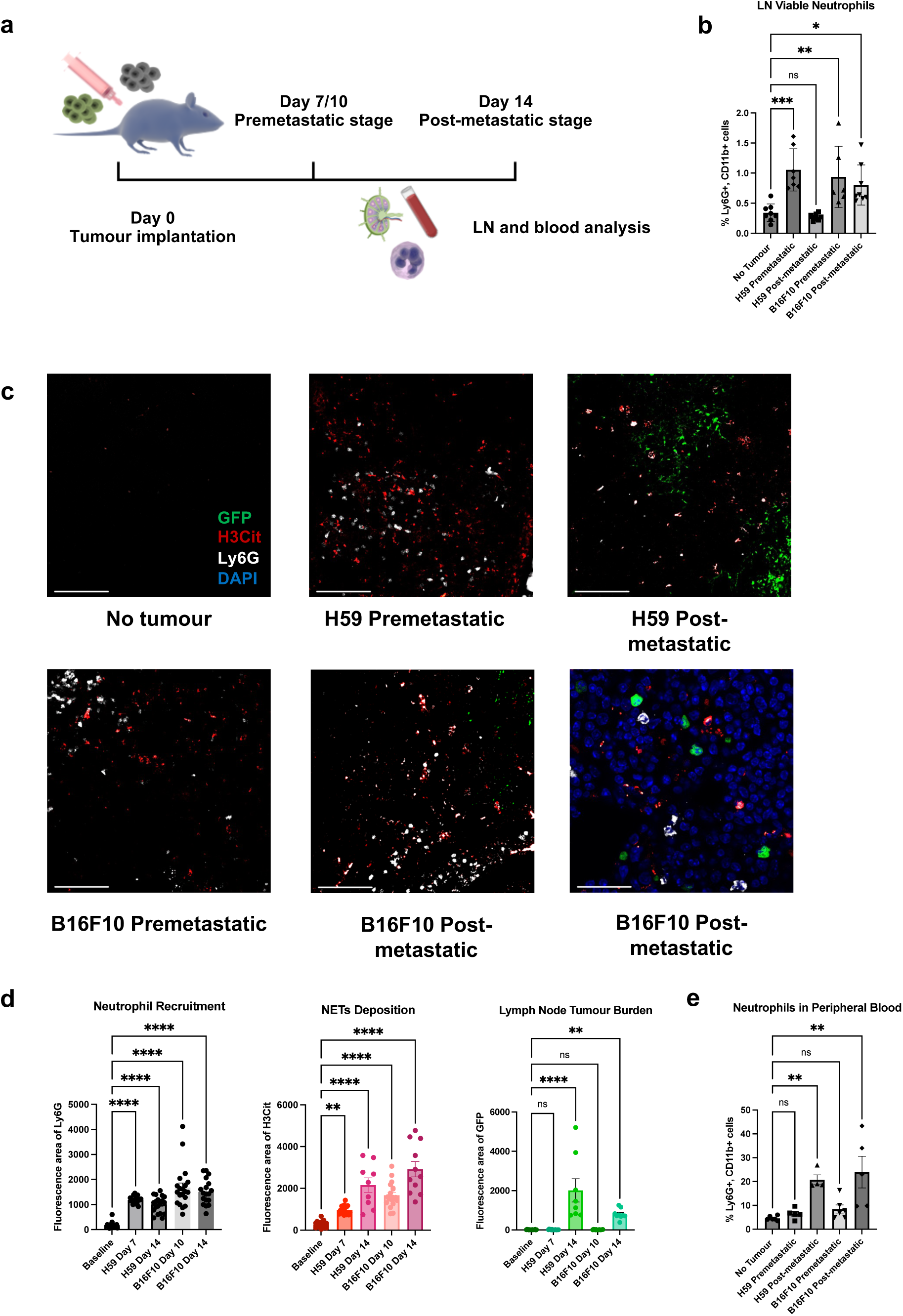
The dynamic of nodal neutrophil recruitment and NETs deposition in mouse model. (a) Schematic illustration of the animal model constructions (b) Percentage of neutrophils within among viable LN cells on a time course by flow cytometry. (c) Representative images of tumour draining LNs on a time course for both H59 lung cancer and B16F10 melanoma. Scale bars represent 100μm. Zoomed details of NETs are shown in bottom right. Scale bars represent 20μm. (d) Quantification of the area of lymphatic neutrophils (Ly6G), NETs (H3Cit) and tumour (GFP) in no tumour, pre- and post-metastatic LNs for both H59 and B16F10. (e) Percentage of neutrophils within blood leukocytes on a time course by flow cytometry. Data shown as mean ± SEM. n = 10; *, P < 0.05; **, P < 0.01; ***, P < 0.005; **** P < 0.001 by One-Way ANOVA.

The plateau in neutrophil number could indicate neutrophil death, possibly through NETosis, the process of NETs formation, occurring during the post metastatic stage. In fact, this was confirmed by immunofluorescence microscopy (Fig. 3c). Compared to non-tumour bearing LNs, we found significantly higher neutrophil infiltration (Ly6G area) and NETs deposition (H3Cit area) in both pre-metastatic and post-metastatic LNs (Fig. 3d). Moreover, we observed that increase of neutrophil percentage in blood (general neutrophilia) only occurred post-LN-metastasis (Fig. 3e), suggesting that the local LN inflammation precedes systemic inflammation.

To confirm our results and remove immunogenic effects of GFP, we performed the same experiment with non-GFP tagged cell lines (Fig. 4a). In addition, the observation that neutrophil accumulation and NETs deposition precede overt lymphatic metastasis suggests that cues from the primary tumour drive the formation of a favourable neutrophil/NET rich environment, which is required for tumour cells to metastasize. Correspondingly, we did not observe an increase in LN neutrophil infiltration or NETs deposition using a low-grade cell line, B16F1, that does not induce LN metastasis in our animal model system (Fig. 4b, c), suggesting this LN NETs formation is also associated with the malignancy of tumour cells.

**Figure 4:**
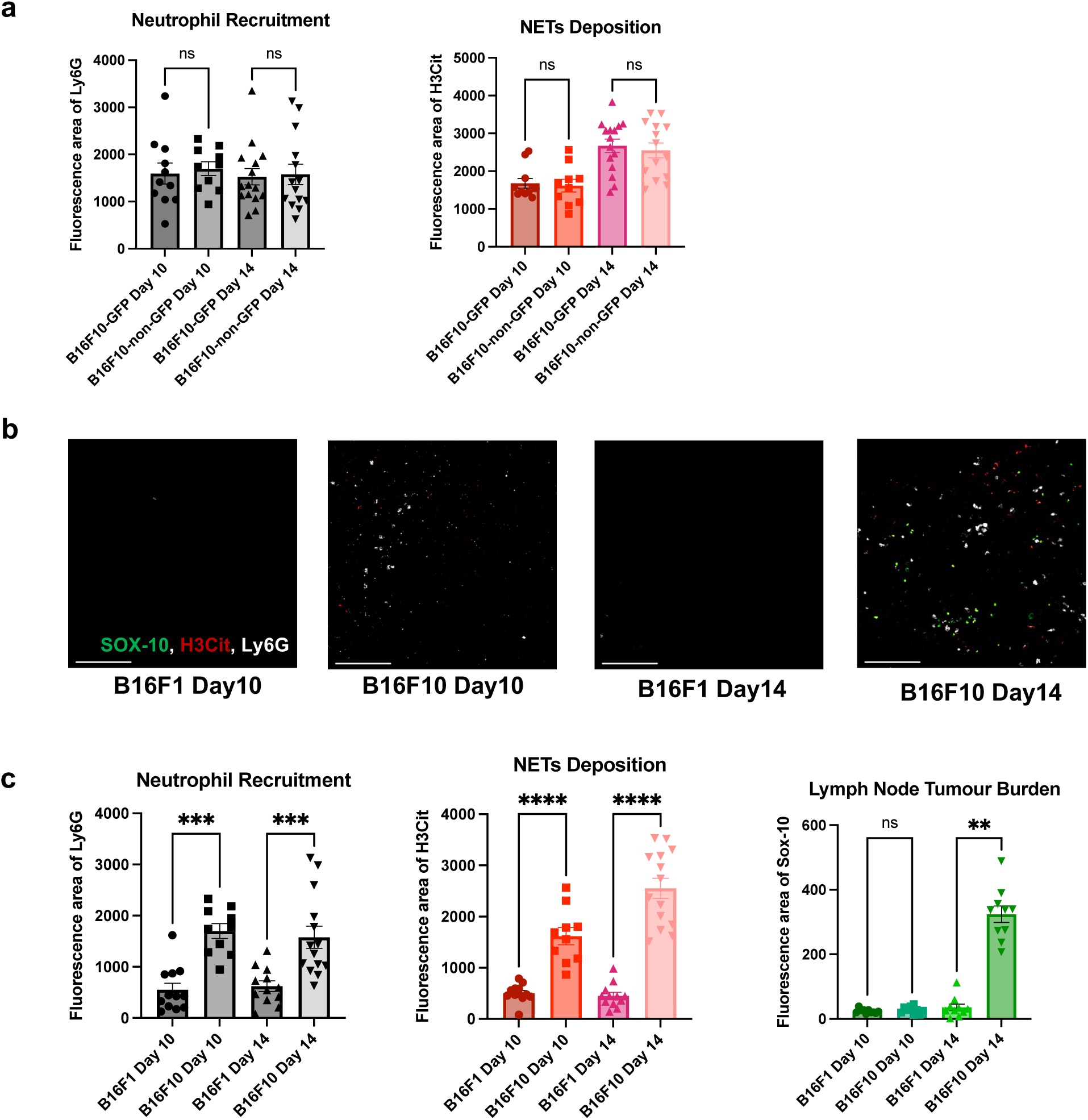
LN neutrophil recruitment and NETs deposition is not associated with the GFP tag but the malignancy of tumour cells. (a) Quantification of the area of lymphatic neutrophils (Ly6G), and NETs (H3Cit) in pre- and post-metastatic nodes for both B16F10 and B16F10-GFP (b) Representative images of tumour draining LNs on a time course for both B16F1 and B16F10 melanoma cells. Scale bars represent 100μm. (c) Quantification of the area of lymphatic neutrophils (Ly6G), NETs (H3Cit) and tumour (Sox-10) in no tumour, pre- and post-metastatic nodes for both B16F1 and B16F10. Data shown as mean ± SEM. n = 10; **, P < 0.01; ***, P < 0.005; **** P < 0.001 by One-Way ANOVA.

### Neutrophil depletion and NETs inhibition abrogates LN metastasis

We hypothesized that NETs play a central role in facilitating LN metastasis. Therefore, we sought to demonstrate this through a number of experiments demonstrating reduced LN metastasis in the absence of NETs (Fig. 5a).

**Figure 5:**
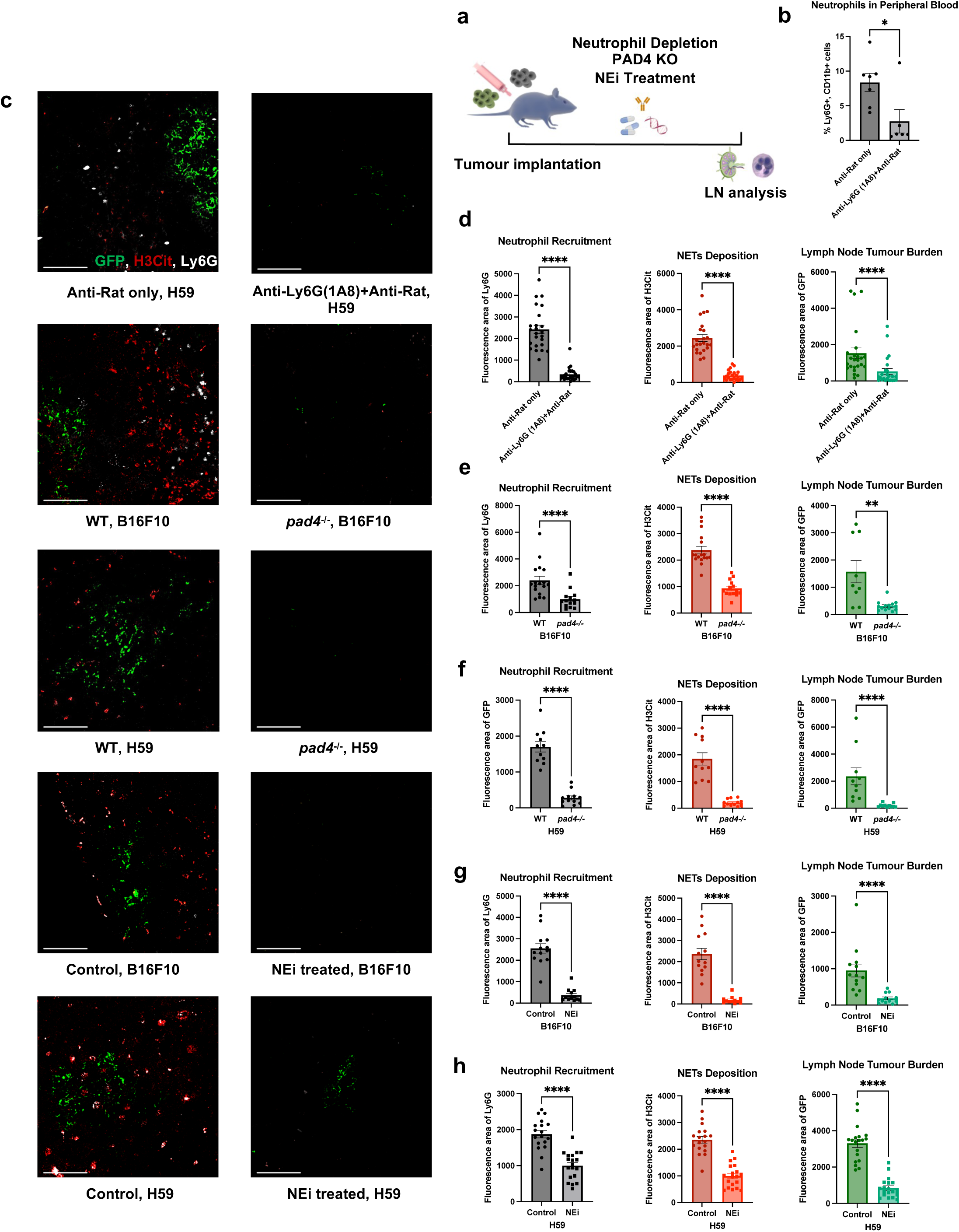
Neutrophil depletion and NETs inhibition abrogates LN metastasis. (a) Schematic illustration of the treatments in animal models. (b) Flow cytometry on blood leukocyte indicates sufficient depletion of neutrophils. (c) Representative images of tumour draining LNs on day 14 post tumour inoculation for both H59 and B16F10, comparing wildtype/no treatment mice to neutrophil depletion, *pad4* knockout and NEi treatment mice. Scale bars represent 100μm. (d)-(h) Quantification of the area of lymphatic neutrophils (Ly6G), NETs (H3Cit) and metastasis (GFP) in day 14 LNs for both H59 and B16F10, comparing wildtype/no treatment mice to neutrophil depletion, pad4 knockout and NEi treatment mice. Data shown as mean ± SEM. n = 10; *, P < 0.05; **, P < 0.01; ***, P < 0.005; **** P < 0.001 by Mann-Whitney t test.

To determine the role of neutrophils in LN metastasis, first, we depleted circulating neutrophils using anti-Ly6G as previously described^57^. We observed a significant reduction in circulating neutrophils on day 14(Fig. 5b) (8.359% vs. 2.755, p=0.023)), suggesting significant depletion. When we assessed the LN, a significantly decrease in neutrophil infiltration, NETs deposition, and metastatic burden was noted (Fig. 5c, d) (neutrophil area 2276 vs 236, NETs area 2109 vs 294, tumour area 1155 vs 229.3, all p<0.0001). The observation that systemic neutrophil depletion could abrogate LN neutrophil recruitment corroborates the association between elevated circulating and lymphatic neutrophils observed earlier in TMAs.

Next, to demonstrate that NETs deposition is required for the effective establishment of LN metastases, we used of Peptidyl arginine deiminase 4 (PAD4) knockout mice (*Pad4*^-/-^), which are unable to form NETs. PAD4 catalyses histone hypercitrullination during NET formation, which is required for histone decondensation^58^. As expected, we observed diminished neutrophil recruitment, NETs formation and metastatic burden in the tumour draining LNs of in *Pad*4^-/-^ compared to wild-type (WT) (Fig. 5c, e and f) (B16F10, neutrophil area 2123 vs. 854.6 p<0.0001, NETs area 2195 vs. 831.6 p<0.0001, tumour area 1061 vs. 302.8 p= 0.0026) (H59, neutrophil area1701 vs. 278.1, NETs area 1849 vs. 208.9, tumour area 1837 vs. 174.7, all p<0.0001). These findings support the essential roles of neutrophils and NETs in the establishment of LN metastases.

The requirement of NETs in development of LN metastasis highlights a potential therapeutic vulnerability. NETs can be pharmacologically targeted with the neutrophil elastase inhibitor Sivelestat (NEi), which prevents NETs formation. Sivelestat and other NEis are an orally bio- available compounds currently under several clinical trials for several autoimmune and respiratory diseases^59, 60^. NEi can inhibits the function of neutrophil elastase, a key regulator and component of NETs^61^. We treated mice daily with NEi treatment and observed a significant decrease in LN NETs deposition and metastatic disease burden (Fig. 5c, g and h). (B16F10, neutrophil area 2444 vs. 243.6, NETs area 2014 vs. 130.1, tumour area 812.9 vs. 130.2, all p<0.0001) (H59, neutrophil area 1701 vs. 278.1, NETs area 1849 vs. 208.9, tumour area 3305 vs. 841.7, all p<0.0001), providing further support for the role of NETs in establishing LN metastasis.

Collectively, these results suggest that recruitment of neutrophils and the subsequent deposition of NETs within LNs are necessary for the establishment of nodal metastasis. Disruption of NETs deposition alone through neutrophil depletion or pharmacologic blockade was sufficient to prevent the formation of clinically detectable LN disease.

### EVs are essential for LN NETs formation and metastasis

In the previous experiment, we demonstrated that lymphatic NETs deposition occurs prior to, and was required for the ingress of neoplastic cells within LNs, it is possible that signalling pathways are originated from the primary tumour and are conducted by tumour derived factors. EVs are actively secreted by nearly all eukaryotic cells and are key players in intercellular communications^62, 63^. Studies have shown that EVs are involved in the formation of a pre-metastatic niche, a fertile microenvironment within target organs favouring neoplastic implantation^64,65,66^. In addition, EVs preferentially accumulate within lymphatic endothelium^52, 53^. Therefore, EVs could provide a blink between the primary tumour and lymphatic inflammatory microenvironment. Interestingly, the expression of genes involved in EV synthesis and secretion are significantly upregulated in oesophageal adenocarcinoma (EAD) patients from The Cancer Genome Atlas Program (TCGA) (Table S2)^67–69^ with nodal diseases. (Fig. 6a) (Rab5a 16.61 vs. 18.96 p=0.0361, PRKD1 0.2598 vs.0.4705 p=0.0240, VAMP7 17.23 vs. 20.91 p= 0.0019). Moreover, high expression of VAMP7 is correlated with poor overall survival in EAD patients (hazard ratio [HR] 2.31, 495 versus 1599 days, p = 0.008) (Fig. 6b).

**Figure 6:**
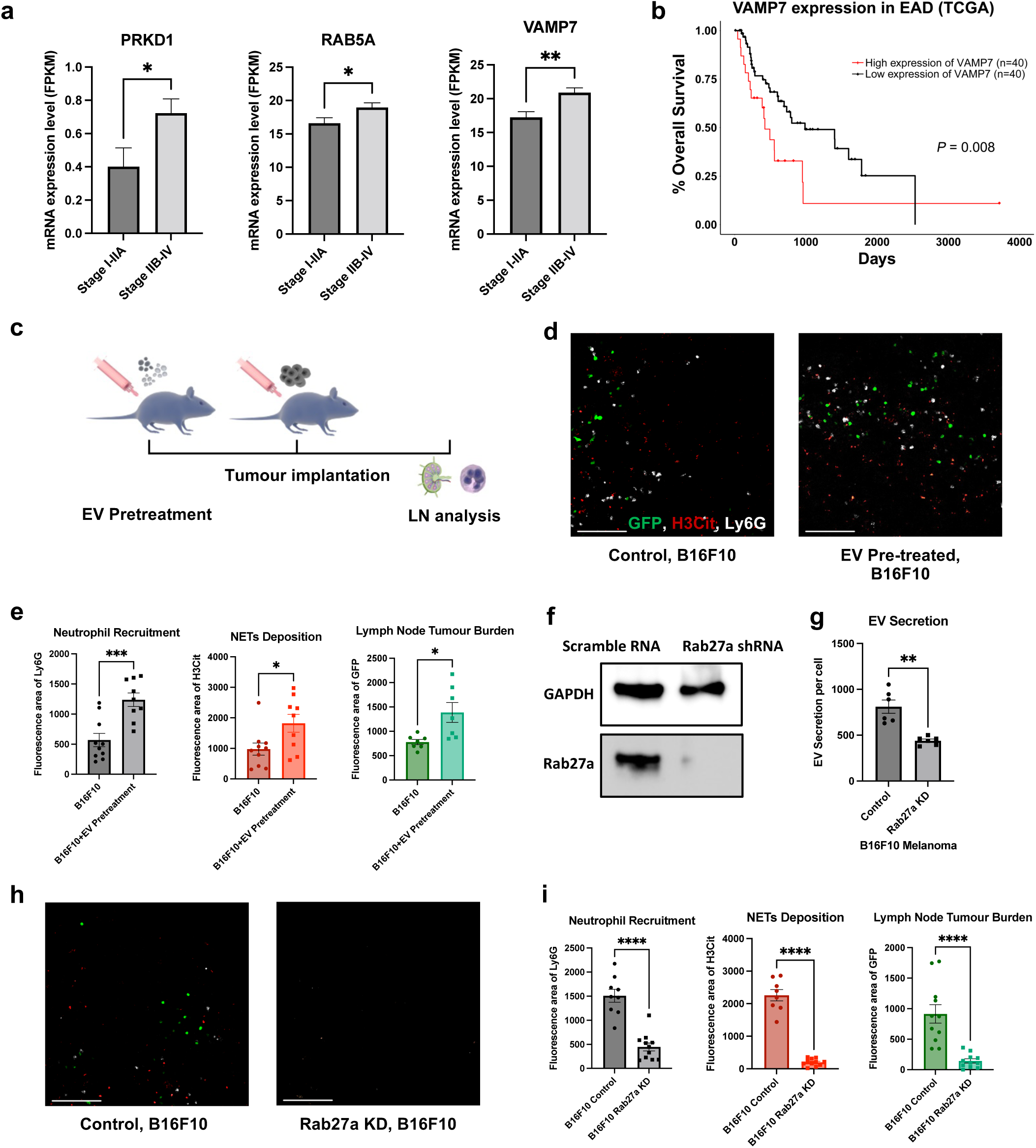
EVs are essential for LNs NETs formation and metastasis. (a) Statistical analysis of TCGA mRNA expression data of EV synthesis related genes in primary esophageal adenocarcinoma (EAD) tumour tissue. n = 15 (Stage I-IIA), n = 65(Stage IIb-IV). (b) Kaplan-Meier survival curves comparing the survival of patients with EAD with low versus high levels of median VAMP7 expression. (c) Schematic illustration of EV pre-treatment in animal models. (d) Representative images of tumour draining LNs on day 14 post tumour inoculation for B16F10, with or without pre-treatment of EVs. Scale bars represent 100μm. (e) Quantification of the area of lymphatic neutrophils (Ly6G), NETs (H3Cit) and metastasis (GFP) in day 14 ;Ns for B16F10, with or without pre-treatment of EVs. (f) Representative Western Blot images indicating the knockdown of *Rab27a* expression in B16F10 cell. (g) Nanoparticle Tracking Analysis (NTA) indicates *Rab27a* knockdown in B16F10 cell leads to decreased EV secretion. (g) Representative images of tumour draining LNs on day 14 post tumour inoculation for B16F10, comparing cells infected with control lentivirus contain scramble RNA or *Rab27a* shRNA. Scale bars represent 100μm. (h) Quantification of the area of lymphatic neutrophils (Ly6G), NETs (H3Cit) and metastasis (GFP) in day 14 LNs for B16F10, comparing cells infected with control lentivirus contain scramble RNA or Rab27a shRNA. Data shown as mean ± SEM. n = 10; *, P < 0.05; **, P < 0.01; ***, P < 0.005; **** P < 0.001 by Mann-Whitney t test.

To validate that EVs from the primary tumour are necessary for lymphatic neutrophil recruitment and NETs deposition, we pre-treated mice with cancer cell derived EVs before tumour inoculation (10μg B16F10 EV per mouse, footpad injection every other day, 3 times) (Fig.6c. and Fig. S1). We found that preexposure to cancer EVs significantly increase nodal neutrophil recruitment and NET deposition (Fig. 6d and e). (Neutrophil area 568.3 vs. 1238 p=0.0006, NETs area 974.1 vs. 1821 p =0.0261, tumour area 779.0 vs. 1387 p=0.0132).

To further confirm the role of EVs, we knocked down (KD) the expression of *Rab27a*, a key gene involved in EV secretion^70^(Fig. 6f). The knockdown in cancer cells approximately halved EV secretion levels (Fig. 6g) (787.0 vs. 450.0 p=0.0022) and resulted in significantly decreased LN neutrophil recruitment, NETs deposition and LN metastasis (Fig. 6h and i) (neutrophil area 1508 vs. 447.1, NETs area 2259 vs. 199.8, tumour area 914.4 vs.144.0, all p<0.0001). Taken together, these data suggest EVs are essential for LN NETs formation and metastasis.

### EV-Lymphatic endothelium interaction results in LN NETs deposition

EVs often exert their biological effects through their uptake into recipient cells and have been shown to preferentially accumulate within lymphatic endothelium^52^. We reconfirmed this finding as the majority of fluorescent EV localised in the cytosol of lymphatic endothelial cells (LECs) in moue LNs (Fig. 7a). In *vitro*, LECs exhibit the same active uptake of EVs (Fig. 7b). Surprisingly, we found LECs dramatically enriched for EVs derived from A549 lung cancer cells compared to EVs from benign bronchial epithelial cells (BEAS-2B), as quantified by the relative area of CFSE-EV to DAPI (Fig. 7c) (0.7436 vs. 0.02310, p<0.0001). Thus, malignant EVs appear to specifically accumulate within lymphatic endothelium.

**Figure 7:**
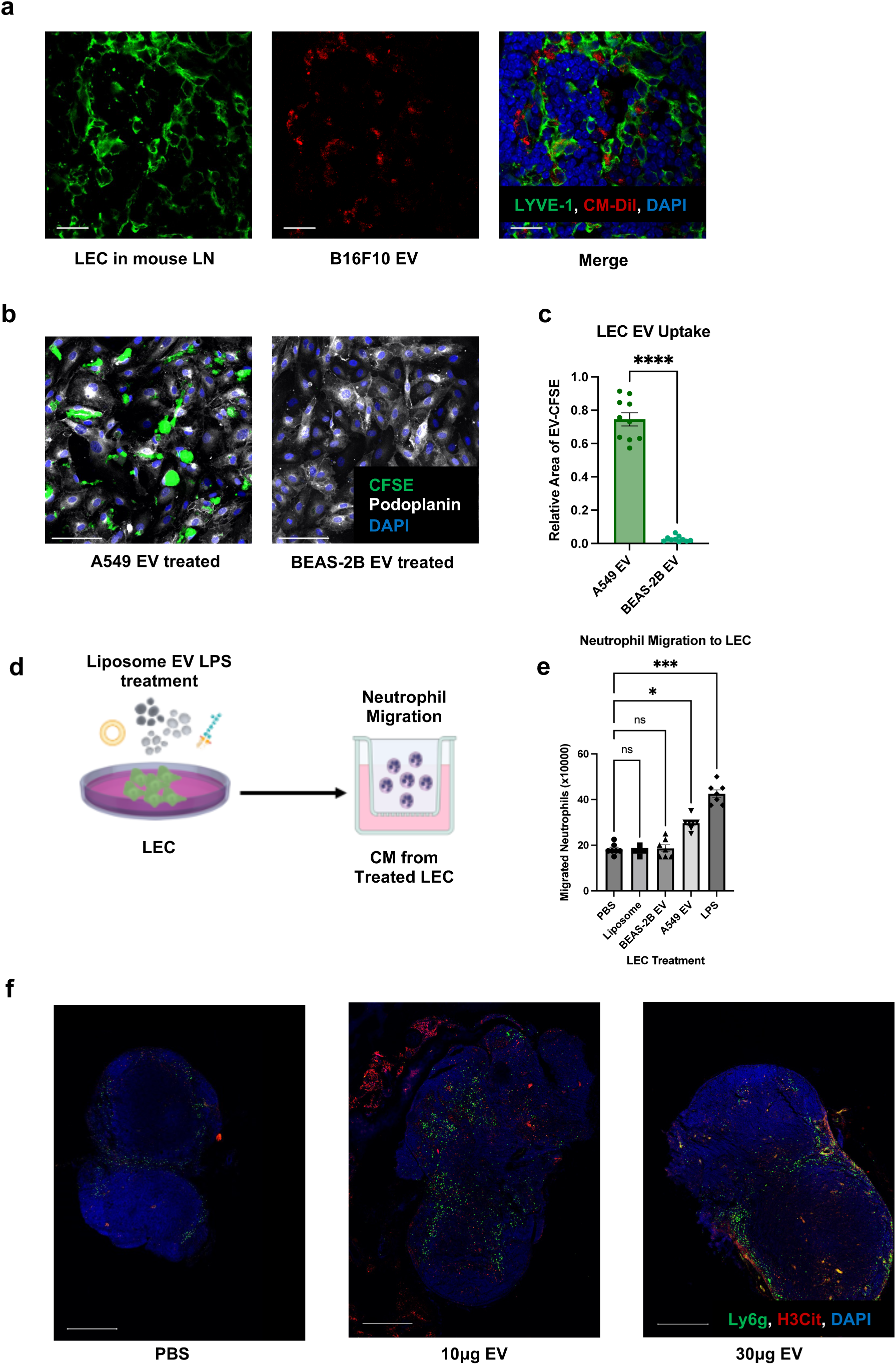
Lymphatic endothelial cells (LECs) are the LN recipient cells of EVs. LECs regulate neutrophil recruitment and NETs formation. (a) Representative images of draining LNs after footpad B16F10 EV injection. Scale bars represent 20μm. (b) Representative images of LEC after A549 and BEAS2B EV treatment. Scale bars represent 100μm. (c) Quantification of the EV uptake (relative fluorescence area of CFSE to DAPI) in LEC. (d)Schematic illustration of the Boyden chamber transwell assay. (e) Boyden chamber transwell assay of neutrophil migration towards conditioned media from treated LECs. (f) Representative images of draining lymph nodes after footpad PBS or different dose of B16F10 EV injection, indicating the subsequent neutrophil recruitment and NETs deposition. Scale bars represent 500μm. Data shown as mean ± SEM. *, P < 0.05; ***, P < 0.005; **** P < 0.001 by Mann-Whitney t test or One-Way ANOVA.

The consequences of LEC EV uptake were subsequently assessed. We performed Boyden chamber experiments to assess the migration of neutrophils towards the conditioned media (CM) from LEC pre-exposed to PBS (vehicle control), liposome (particle control), BEAS-2B EVs (benign EV control), A549 EVs and LPS (positive control) (Fig. 7d). We found that LECs treated with cancer derived EVs could significantly induce neutrophil transwell migration, while LECs treated with EVs from benign cells or liposomes could not (Fig. 7e) (24.93 vs 18.64/17.5 x10^6^ neutrophils, p=0.0489). The finding of EV mediated neutrophil accumulation and NET deposition was subsequently confirmed *in vivo*. B16F10-derived EV were injected into the footpad of non-tumour bearing mice. Interestingly, EV administration alone is sufficient to drastically increase LN neutrophil infiltration as well as the deposition of NETs in a dose-dependently manner (Fig 7f and Fig. S1a) (PBS VS 10μg EV: neutrophil area 12.95 VS.28.14 P=0.0026, NETs area 15.57 vs. 18.00 not significant; PBS VS 30μg EV: neutrophil area 12.95 vs. 26.00 p=0.0050, NETs area 15.57 vs. 27.78 p=0.0093).

### Lymphatic secretion of CXCL8/2 induces LN Neutrophil infiltration and NETs formation

Having demonstrated that tumour EVs are sufficient to induce lymphatic neutrophil migration and NETs deposition, we next sought to elucidate the mechanism by which this occurs. First, we collected conditioned media (CM) from LECs that were co-cultured with EVs from A549 lung cancer cells or BEAS-2B bronchial epithelial cells (Fig. 8a). We then performed a multiplex ELISA targeting common neutrophil chemokines. Interestingly, LEC production of CXCL8 was radically increased after A549 cancer cell derived EV treatment (Fig. 8b) (1865 vs. 4447 pg, p=0.0004). Moreover, A549 cells themselves also secreted more CXCL8 compared to BEAS-2B (91652 vs. 16316 pg, p <0.0001). We also performed immunofluorescence to detect LEC derived CXCL8 and observed a significant increase in CXCL8 synthesis in LECs following A549-derived EV treatment (Fig. 8c, d), quantified by relative area of CXCL8 to DAPI (5.500 vs. 20.40 p= 0.0003). In addition, we also demonstrated the LEC expression of CXCL8 in surgical lymph node samples from lung adenocarcinoma patients (fig 8e).

**Figure 8:**
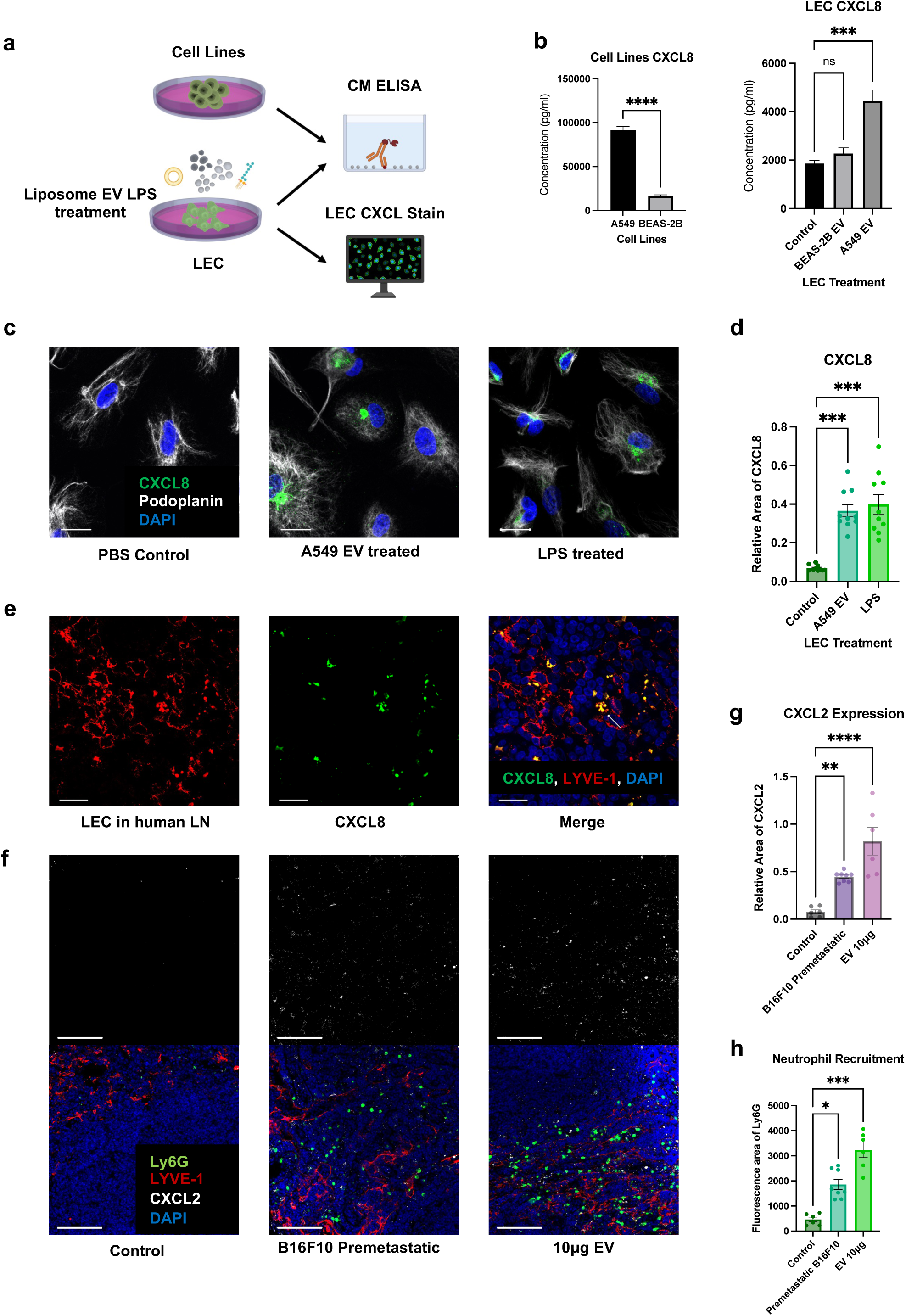
LECs secrete neutrophil chemoattractants and NETs inducers upon EV uptake. (a)schematic illustration of the ELISA and LEC CXCL8 stain. (b) ELISA of CXCL8 level in conditioned media (CM) of A549, BEAS-2B and treated LEC. (c) Representative images of LEC expressing CXCL8 different kinds of treatments. Scale bars represent 20μm. (d) Quantification of CXCL8 expression in LECs (relative fluorescence area of CXCL8 to DAPI). (e) Representative images of LECs expressing CXCL8 in positive nodes of lung cancer patients. Scale bars represent 20μm. (f) Representative images of LEC expressing CXCL2 and recruiting neutrophils mouse LNs. Scale bars represent 100μm. (g) and (h) Quantification of CXCL2 expression (relative fluorescence area of CXCL2 to LYVE-1) in LEC and neutrophil recruitment (Ly6G fluorescence area) in mouse LNs. n = 5. Data shown as mean ± SEM. *, P < 0.05; **, P < 0.01; ***, P < 0.005; **** P < 0.001 by Mann-Whitney t test, One-Way ANOVA or Kruskal-Wallis test.

Mice do not express CXCL8 but do express its analogue CXCL2^71^. We found CXCL2 expression colocalised well with LECs and was in proximity to LN neutrophils (Fig. 8f). The level of CXCL2 was also extensively increased in pre-metastatic condition or following EV injection (Fig. 8g), quantified by relative area of CXCL2 to LYVE-1 (control vs. B16F10 premetastatic 0.07592 vs. 0.4462, p= 0.0058; control vs. EV 10μg 0.07592 vs. 0.8199, p <0.0001), which was accompanied by an increase in the number of neutrophils infiltrated into LNs, quantified by area of Ly6G (Fig. 8h) (control vs. premetastatic B16F10 3.500 vs. 10.88, p= 0.0420; control vs. EV 10μg 3.500 vs. 17.00, p=0.0002).

In addition to a common chemoattractant of neutrophils, CXCL8 is also a potent NETs inducer^72^. To consolidate the conclusion that cancer derived EVs can induce LEC to secrete CXCL8, thus resulting in NETs formation, we exposed healthy control-derived neutrophils to CM from LECs that were treated with A549 cancer cell derived EV with or without a CXCL8 blocking antibody (Fig. 9a). We found that CM from A549 EV treated LECs significantly increased the percentage of neutrophils that were undergoing NETs formation and CXCL8 blockade abrogated this phenotype (Fig. 9b and c) (control vs. A549 EV 7.415% vs. 79.63%, p<0.0001; control vs. A549 EV+CXCL8 Ab 7.415% vs. 24.64% ns; A549 EV vs. A549 EV+CXCL8 Ab 79.63% vs. 24.64%, p<0.0001). Furthermore, since EVs can directly lead to the release of NETs^55^, we simultaneously exposed GEA patient or healthy control derived neutrophils to A549-derived EVs and we found that GEA patient derived neutrophils are more likely to form NETs (Fig 9 d and e)(39.71 vs. 23.36 p= 0.0040). The increased propensity of NETosis in response to EVs in cancer patient indicates the profundity of neutrophil preexposure to other tumour derived molecules in cancer patients and the potential synergy of EVs and cytokines^73^.

**Figure 9:**
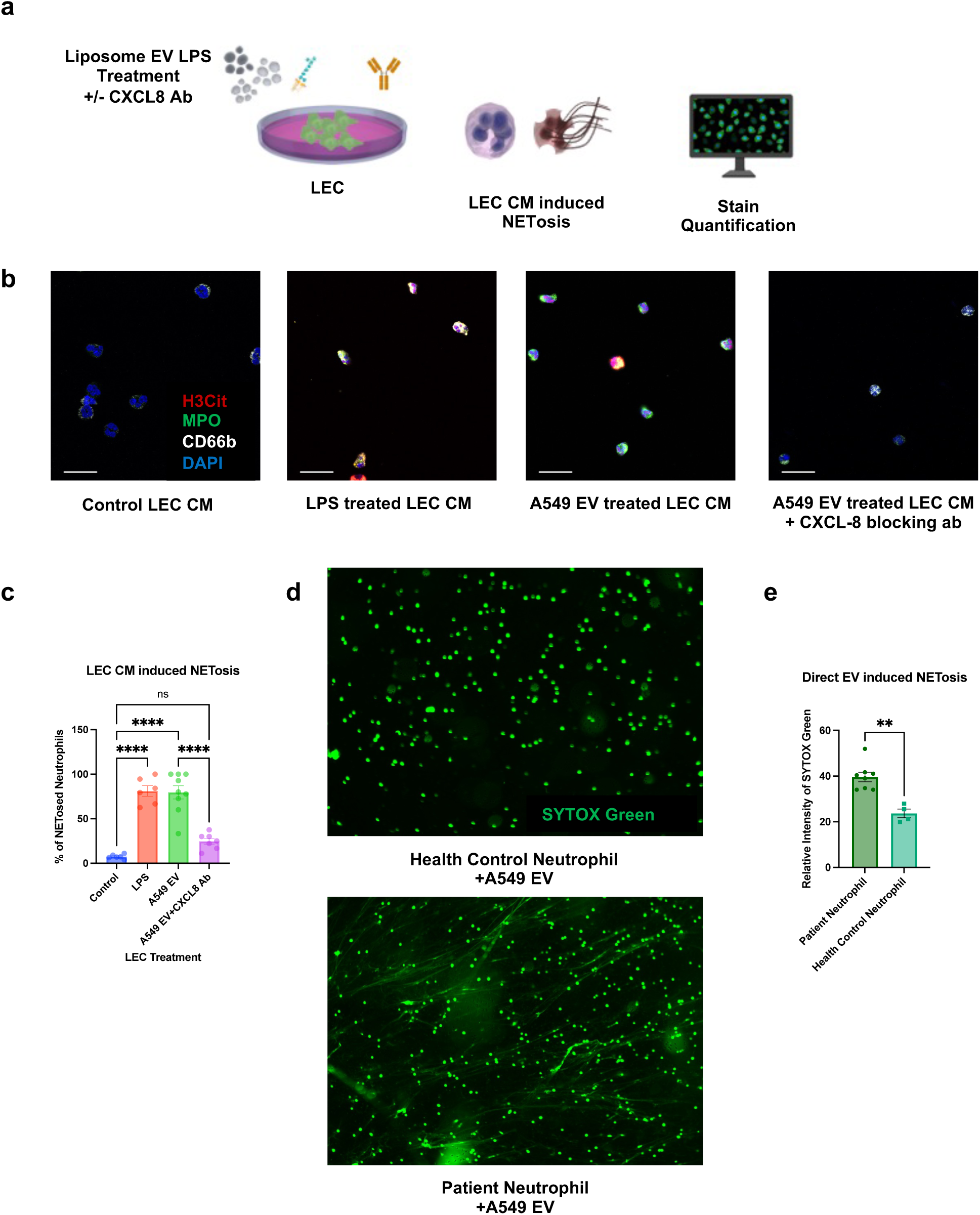
Lymphatic CXCL8 secretion upon EV uptake leads to NETs formation. (a)Schematic illustration of the in vitro NETosis assay induced by CM of treated LECs. (b) Representative images of neutrophils treated with CM from treated LECs. Scale bars represent 20μm. (c) Quantification of LEC CM induced NETosis (% of H3Cit positive neutrophils). (d) Representative images of indicating the different proneness to form NETs for neutrophils from health control and GEA patients after A549 EV treatment. Scale bars represent 100μm. (d) Quantification of propensity of neutrophil to form NETs induced by EV (Relative fluorescence intensity of SYTOX Green treated to untreated neutrophil) from health control and GEA patients). Data shown as mean ± SEM.**, P < 0.01; ****, P < 0.001 by Kruskal-Wallis test or Mann-Whitney t test.

## Discussion

LN metastasis is nearly ubiquitous among solid tumours and portend poor survival^74^. Treating regional LNs via resection forms the basis for modern surgical oncologic treatment approaches. Regional LNs are removed and often trigger the implementation of systemic therapies when they harbour cancer^75^. The commonality of LN metastasis among cancers presents a broadly applicable opportunity for treatment. Despite this, little is understood about LN metastasis actually forms. In particular, the link between the primary tumour site and regional LN remains poorly described.

In the current study we set out to demonstrate that similar to conditions of systemic inflammation, a regional inflammatory state can promote the development of LN metastasis. We demonstrated that lymphatic neutrophil accumulation and NETs deposition occurs within the tumour LNs of GEA patients. Furthermore, we reveal that the level of lymphatic NETs is an indicator of poor prognosis among these patients. Interestingly, while NETs levels were highest in LNs that had been infiltrated with cancer (N+met), even negative regional nodes (N+neg) demonstrated elevated NETs compared to LNs harvested from patients without nodal metastasis (N0). This finding suggests that regional lymphatic neutrophil accumulation and NETs deposition can occur without the presence of neoplastic cells. Since NETs levels were highest in N+met, we surmised that lymphatic neutrophils and NETs present a favourable environment for tumour implantation. As such, perhaps they should represent a treatment target in and of themselves.

We show that regional lymphatic inflammation caused by neutrophils results from inflammatory mediators, in the form of EVs, originating from within the primary tumour. This distinct premetastatic phase, within LNs, is characterized by lymphatic neutrophil infiltration and subsequent deposition of NETs. Our data, on multiple cancer types, confirm that LN metastasis together with the regional LN inflammation we discovered is a universal behaviour of cancers, and it is induced by environmental cues from the primary tumour. Correspondingly, in the study by Wang *et al*, the authors highlight tumour associated neutrophils (TAN) within early gastric cancer resection specimens as independent risk factors for LN metastasis^76^. Similarly, Hiramatsu *et al* demonstrate a similar association between the number of neutrophil number in the tumour draining LN and patient outcome^77^. In the data presented here, regional lymphatic inflammation by neutrophils appears to have increased as a result of inflammatory mediators in the form of EVs possibly originating from within the primary tumour. The observation of a distinct pre-metastatic phase has important implications when considering the pathologic staging of neoplasia. LN staging in contemporary clinical practice is binary; nodes are either infiltrated with cancer or not. This study, along with previous work on the prognostic profundity of looking art immune compartment in pathological samples, suggests LN infiltration represents a spectrum of disease^78–80^.

EVs derived from the primary tumour play a critical role in establishing a pre-metastatic niche within LNs. Tumour derived EVs have been shown to rapidly disseminate through the lymphatic system and into regional LNs, particularly during infection^53^. The observation that EVs can be rapidly taken up by LECs, transported to the lumen of the vessel, and concentrated rapidly within LNs has led to the suggestion that they represent a means of information exchange that can precede the arrival of migrating cells. We posit that an analogous mechanism is taking place between the primary tumour site and regional lymphatics. Supportingly, lymphatic fluid from patients with cutaneous melanoma are enriched with EVs derived from their tumours^52^. Therefore, an analysis of lymphatic composition may permit the stratification of patients with early as opposed to advanced disease.

Neutrophil depletion abolished both the deposition of presumed NETs and abrogated the formation of nodal metastasis all together^41, 44, 45^. We and others have revealed a pivotal role for neutrophils in the development of metastasis^16, 34^. In the current study, we observed widespread NETs in LNs, which led to the formation of an environment permissive to tumour outgrowth. Indeed, NETs have been shown to play a pro-tumorigenic role through direct effects on tumour cell proliferation, invasion, and escape from dormancy^44–46, 81^, Furthermore, NETs themselves have been shown to induce local immune suppression through the extracellular elaboration of NET associated PD-L1^82^. Thus, it is feasible to surmise that NETs supported tumour outgrowth through a variety of mechanisms, including but not limited to, tumour cell sequestration, enhanced adhesion, migration, proliferation, and local immune suppression. We confirmed the in the process of LN metastasis through the effect of both PAD4 inhibition and NE inhibition on tumour outgrowth.

EVs were sufficient to induce nodal neutrophil ingress in *vivo* and were necessary for the formation of metastasis. Similar findings have been reported for macrophage recruitment^83^, lymphangiogenesis^84^, and immune suppression^13^ within LN microenvironment. Indeed, EVs serve as mediators of communication between primary tumour cells and LN cells, resulting in premetastatic changes that create a permissive condition for subsequent tumour cell colonisation. Our work highlights the essential role of EVs in mediating this NET-induced LN metastasis. Moreover, while the process of NET formation has been reported to be regulated by both cytokines, such as CXCL8 and G-CSF^72, 73^, in addition to EVs^55^, we are the first to establish a link between the regulation of the CXCL8, EVs and NETs formation.

EVs induced the production of CXCL8 leading to neutrophil migration and NET deposition both in *vitro* and in *vivo*. This was confirmed through CXCL8 inhibition which diminished much of this phenotype. CXCL8 is a pro-inflammatory cytokine that predominantly activates and recruits neutrophils^85^. Earlier work has characterised the role of CXCL8 as a NET inducer and aggravator in chronic inflammatory diseases. Future works is needed to access the synergy between tumour-derived soluble factors and EVs and investigate the plasticity of neutrophil function in the milieu of cancer from a cancer secretome mindset.

In summary, the data presented sets a scene wherein tumour derived EVs accumulate within lymphatic endothelium. This results in a local chemotactic gradient, involving CXCL8, that promotes neutrophil influx and NET deposition. Furthermore, locally infiltrated neoplastic cells may themselves attract neutrophil ingress and induce NET release by releasing more CXCL8 and EVs. The NETs deposition results in a local microenvironment that is permissive and favourable to tumour outgrowth. This was the case in the murine models employed. In addition, there was a clear association between lymphatic NETs deposition and reduced survival in gastroesophageal cancer patients. The finding that this may occur across malignant types may highlight NETs as a potential therapeutic target based not on histology but host inflammatory status instead. Given that no treatments to date target host inflammatory status, the results of this study are particularly germane. We highlighted that NETs represent an attractive therapeutic target as they are amenable to pharmacologic inhibition and appear to be involved throughout the metastatic cascade, as well as the therapeutic potentials of targeting EVs and CXCL8. Future works further investigating clinical efficiency of NETs targeting agents in treating LN metastasis should lead to major advances in the management of cancer patients.

## Materials and Methods

### Patients

This study investigated surgical lymph node of 175 patients with stage I to IV gastroesophageal adenocarcinoma (Table S1) who were treated at the McGill University Health Centre (MUHC), Montréal, Canada. All the studies were conducted according to the relevant regulatory standards, upon approval by the Research Ethics Board Office at McGill University.

### TMA construction, immunofluorescence (IF) staining and analysis

Tissue microarrays (TMAs) were constructed from the surgical blocks used for pathologic evaluation. Three pathologists constructed 8 TMA blocks using 1 mm cores punched from formalin-fixed and paraffin-embedded tissue blocks. After sectioning of TMA blocks and IHC staining, a pathologist verified that each core contained the lymph node tissue of interest and whether there was LN metastasis. TMA slides were stained at the histopathology core of Research Institute of McGill University Health Centre (RI-MUHC) using primary antibodies including anti-NE (MAB91671100; R&D Systems), anti-CD66b (392902; BioLegend), anti-H3Cit (ab5103; abcam) and anti-cytokeretin 7(ab9021; abcam). To visualize these antibodies, OPAL kit (PerkinElmer) fluorophores were used and prepared according to the manufacturer’s protocols. The slides were counterstained with DAPI (D21490; Invitrogen) and cover slipped. TMA slides were then digitalized at 20× magnification using Axio Scan Z.1. whole slide scanner (Zeiss). HALO image analysis software (Indica Labs) was used to quantify the number of CD66b^+^NE^+^ positive cells as neutrophil and the area covered by NETs(H3Cit).

### Quantification of neutrophil and lymphocyte counts in peripheral blood

Circulating NLR of patients was calculated from clinical laboratory blood tests performed on the date of diagnosis. A cut off NLR value of 4 was used to assign patients to into low or high groups. LN NLR was determined via IF staining of TMAs as described before.

### Cell Lines

B16-F10, H59, A549 and BEAS-2B cells were all maintained in corresponding media suggested by American Type Culture Collection (ATCC) with 10% heat inactivated FBS. Cells used for in vitro EV production were maintained in media with 10% heat inactivated FBS depleted of bovine EVs by ultracentrifugation at 100,000g for 14 hours.

### Animals

C57BL/6 (Charles River Laboratories) and PAD4 knockout (PAD4^−/−^; gift of Alan Tsung, The Ohio State University Comprehensive Cancer Centre, Columbus, Ohio, USA) mice were used for all experiments at 7–10 weeks old. 250k B16F10-GFP, H59-GFP, B16F10, B16F1, B16F10-scramble RNA, or B16F10-Rab27 KD cells were injected subcutaneously into the flank of these mice, and tumour growth was monitored twice a week using a calliper. Blood was collected by saphenous vein bleeding. All experiments were performed with the Veterinary Authority of the Institutional Animal Care and Use Committee at the RI-MUHC under protocol no. 7469 and 8210. For the neutrophil depletion, experimental group mice received intravenous injection of 100μg anti-Ly6G (1A8, BE0075-1; BioXCell) on day 0, 2, 6, 8, 10, 12 post tumour inoculation, and 100μg of anti-Rat-kappa light chain (MAR18.5, BE0122; BioXCell) on day 1, 5, 7, 11, 13 post tumour inoculation. Control group only received anti-Rat-kappa light chain injection. For the knockout experiments, PAD4–/– mice were compared to age and gender matched C57BL/6 mice. For the NEi treatment, experimental group mice received daily gavage of 2.2 mg/kg of neutrophil elastase inhibitor Sivelestat (NEi, ab146184; abcam) resuspended in saline. Control mice received saline gavage.

### Flow cytometry

Mouse blood were prepared by lysing red blood cell in BD Pharm Lyse lysing solution (555899; BD Pharm) and washed in PBS. Mouse LN were minced and filtered through 40μm cell strainers (087711; Fisher Scientific) and washed in PBS. Cell suspensions were blocked with Fc-block (553141; BD) and incubated with the following primary antibodies: Viability Dye Eflour780 (65-0865-14; eBioscience), anti-CD11b-BV510(101263; BioLegend) and anti-Ly6G-AF647(127610; Biolegend). Data acquired on a BD LSRFortessa™ Cell Analyzer cytometer was analyzed using FlowJo software (Tree Star).

### Lymph node immunofluorescence staining, imaging, and analysis

Mouse Ipsilateral inguinal lymph node of the tumour side was dissected after sacrificing and fixed in 1% paraformaldehyde, dehydrated in 30% sucrose, and embedded in Optimal Cutting Temperature (OCT) compound. 15 μm OCT tissue sections were stained with primary antibodies including: anti-H3Cit (ab5103; abcam), anti-Ly6G-AF647 (127610; Biolegend), anti-SOX-10-AF488 (NBP2-59621AF488; Novus Bio),anti-LYVE-1 (AF2125; R&D Systems), anti-Ly6G-FITC (11-9668-80; Biolegend), anti-MIP2/CXCL2 (500-P130; PeproTech) and secondary antibodies including: goat-anti-rabbit-AF568 (A11011; Invitrogen), donkey-anti-goat-AF488 (A110555; Invitrogen), donkey-anti-goat-AF568 (A11057; Invitrogen), donkey-anti-rabbit-AF647 (A-31573; Invitrogen).

Human lymph node block from consented lung adenocarcinoma patients was retrieved from Department of Pathology of MUHC. 4μm paraffin tissue slides were de-paraffined, antigen-retrieved in pH6 sodium citrate buffer and stain with primary antibodies including: anti-LYVE-1 (AF2089; R&D System), anti-CXCL8 (MAB208; R&D System) and secondary antibody including: donkey-anti-goat-AF568 (A11057; Invitrogen) and donkey-anti-mouse-AF488(A-21202; Invitrogen).

The slides were counterstained with DAPI (D21490; Invitrogen) and cover slipped. Negative controls were stained with secondary antibody only or isotypes conjugated to the same fluorophore. The slides were imaged with LSM 780 confocal microscope at the molecular imaging core of RI-MUHC. ImageJ Software (NIH) was used to analyse the fluorescence area of GFP, Ly6G, H3Cit, SOX-10 and the relative fluorescence area of CXCL2 to LYVE-1.

### EV isolation labelling and mouse injection

Cells were cultured in media supplemented with 10% EV-depleted FBS. Supernatant fractions collected from 48–72 h cell cultures were pelleted by centrifugation at 300g for 10 min. The supernatant was centrifuged at 20,000g for 20 min. EVs were then harvested by centrifugation at 100,000g for 70 min. The EV pellet was resuspended in 20 ml of PBS and collected by ultracentrifugation at 100,000g for 70 min. For fluorescence labelled EVs, supernatant was concentrated using Amicon Ultra-15 Centrifugal Filter Units with 100 KDa filter size, incubated with 1.0 mM CM-DiI or 50 µM CFSE for 2 hours before this first 100,000g centrifugation.

For the EV pre-treatment experiment (Fig. c, d), 10 µg EV in 30 µL PBS was injected into the ipsilateral footpad every other day before tumour inoculation. For the EV location experiments, 10 µg CM-Dil labelled EV were injected 24 hours before sacrificing mice and LN dissection. For the EV treatment in no-tumour bearing mice, 10-30 µg EV in 30 µL PBS was injected into the ipsilateral footpad every other day before sacrificing.

### Lentiviral infection and screening procedure

B16F10 cells were transfected with Rab27a-mouse shRNA and scramble RNA lentiviral particles purchased from Origene according to the manufacturer’s instructions. Briefly, B16F10 cells were plated in 24-well plate at 50,000 cells per well (day zero). At day 1, cells were infected with lentiviral vectors in the presence of 8 μg/ml polybrene. Media was changed at day3 and from day 4, cells were treated with 0.5 μg/ml puromycin for 10 days to generate the stable knockdown cell line. The sequence of the Rab27a shRNA is CTGGATAAGCCAGCTACAGATGCACGCGT.

### Western Blot

25 μg of EVs or cell lysates was mixed with Laemmli SDS sample buffer (Bio-Rad), incubated 10 min at 95°C, and cooled to 4°C. Electrophoresis was performed on Mini-PROTEAN TGX Gels

(Bio-Rad). Proteins were transferred to a nitrocellulose membrane (Bio-Rad). Blocking was performed 30min at room temperature (RT) in 5% Tris-buffered saline (TBS) milk, primary in 5% TBS bovine serum albumin (BSA), overnight at 4°C (shaking), and secondary (HRP-conjugated;Bio-Rad) in 5% TBS BSA, 30min at RT (shaking). The following primary antibodies were used: anti-ITGA6 (1:250, 3750; Cell Signaling), anti-Alix (1:250, 2171; Cell Signaling), anti-BiP (1:500, 3183; Cell. Signaling), anti-TSG101 (1:500, ab125011; abcam), anti-GAPDH (1:1000, MA5-15738; Invitrogen) and anti-Rab27a (1:500, 69295; Cell Signaling).

### Nanosight analysis of EVs

EV density and size distribution were assessed by nanoparticle tracking analysis (NanoSight; Malvern Instruments). 5 μl of isolated EVs was diluted in 500 μl of PBS to achieve a uniform particle distribution that was analyzed in three to five sequential measurements at 37°C. For the Rab27a KD experiments, 500ul of FBS-free cell conditioned media from control and Rab27a KD cell lines were used.

### Transmission electron microscope (TEM) analysis of EVs

Negative stain was performed in the following way: 5 μl sample solution was adsorbed to a glow-discharged carbon-coated copper grid (Canemco & Marivac), washed with deionized water, and stained with 5 μl 2% uranyl acetate. The samples were imaged at RT using Tecnai Spirit electron microscope at the Facility for Electron Microscopy Research of McGill University.

### LEC culture and immunofluorescence

LEC were purchased and cultured MV2 media (C-22022) from in PromoCell on Collagen I coated tissue culture vessels and were treated with 10 μg/ml EV or 5 μg/ml Lipopolysaccharides (LPS, Sigma-Aldrich) for 24 hours. The LECs monolayer was washed in PBS after cocultures was done. After 15 min of fixation n 2% paraformaldehyde, RT, LEC were stained with anti-Podoplanin-AF647 and anti-CXCL8-AF488 (IC208G; R&D System). Negative controls were stained with isotypes conjugated to the same fluorophore. The slides were imaged with LSM 780 confocal microscope at the molecular imaging core of RI-MUHC. ImageJ Software (NIH) was used to analyse the fluorescence area of CFSE and the relative fluorescence area of CXCL8 to DAPI.

### Human peripheral blood neutrophil isolation

Human neutrophils were isolated as previous described^44^. Briefly, blood was diluted in PBS and layered over Lymphocyte Separation Media (Wisent Bioproducts). After centrifugation at 800 g for 30 min at RT, the pellet containing neutrophils and red blood cells (RBCs) was collected and resuspended in 3% Dextran (Wisent) and left at RT to sediment RBCs. After 30min, the supernatant is collected and centrifuged at 450 g at 4°C for 5 min. Remaining RBCs in the neutrophil rich pellet are then lysed using BD Pharm Lyse Lysing buffer. The obtained pellet is then washed and resuspended in cold RPMI media (Wisent Bioproducts. Purity and viability of the obtained neutrophils were verified through Methylene blue (Stem cell Technologies) and Trypan blue (Wisent) staining respectively.

### Boyden Chamber Neutrophil migration assay

LEC were treated with 10 μg/ml A549/BEAS-2B EV, 5 μg/ml LPS, 100nm Liposome control (Encapsula Nanoscience) of same number of particles (determined by NTA) or PBS of the same volume in MV2 for 24 hours. The next day, fresh MV2 media was changed. 72 hours later. The conditioned media was collected, centrifuged at 450g at 4°C for 5min to pellet down cells. The supernatant was then transfer to the bottom chamber of the transwell. 1 million neutrophil was layered in the top chamber and leave for migration. 24 hours later, migrated neutrophil in the bottom chamber was resuspended and count by Methylene blue stain. This assay was repeated 3 times with triplicates.

### ELISA of CXCL8

The A549/BEAS-2B cell lines were incubated in corresponding media with 0.5% FCS for 72 hours to collect cell conditioned media (CM). The LEC CM were collect in the same was as the boyden chamber experiments. CXCL8 amount in the CM were incubated with fluorescence multiplex ELISA kit from RayBioTech and quantified using Q-analyser from the same manufacturer.

### LEC CM induced NETs formation

LEC CM was collected in the same way as in the boyden chamber experiments and transferred to Nunc Lab-Tek II Chamber Slide (Thermo Scientific), 200k neutrophils were added to the CM and NETs formation was performed at 37°C. After 24 hours, the supernatant was discarded and the slides were gentle washes, fixed with 4% PFA for 15min and stained with DAPI, anti-MPO-FITC (ab11729; abcam), anti-H3Cit (ab5103; abcam), anti-CD66b-APC (130-122-966; Miltenyl Biotech) and then goat-anti-rabbit-AF568 (A11011; Invitrogen). The slides were imaged with LSM 780 confocal microscope at the molecular imaging core of RI-MUHC. Percentage of NETosing neutrophils (H3Cit positive neutrophils) were quantified.

### EV induced NETs formation in health control and GEA patients

Health control or GEA patient neutrophils were isolated as mentioned before. Every 200k neutrophil were treated with 10 μg/ml A549 EV and incubated at 37°C for 4 hours. After 4 hours, SYTOX Green (Invitrogen) was added to a final dilution of 1:1000 and fluorescence intensity were acquired Tecan Infinity F200. Proneness to NETose was quantified as the relative fluorescence intensity of SYTOX Green of EV treated wells to untreated wells. Representative images were acquired using EVOS cell imaging systems (Thermo Scientific).

### Statistical analysis

Data were expressed as the mean ± SEM. Data were analyzed using Prism software (GraphPad Software, Inc.). Statistical significance between two groups was assessed using a Mann Whitney t test. When more than two value sets were compared, we used one-way ANOVA, Kruskal-Wallis test or Brown-Forsythe ANOVA test. P < 0.05, P < 0.01, P < 0.001, or P < 0.0001 was considered statistically significant.

The Kaplan-Meier curves in Fig.2c were generated using Prism software and the Kaplan-Meier curves in Fig. 5a were generated using RStudio (Version 3.6.3)^86^. Higher and lower expression levels were stratified based on median expression levels. Statistical analysis was performed using log-rank tests, and HRs were calculated.

---

**Figure S1:**
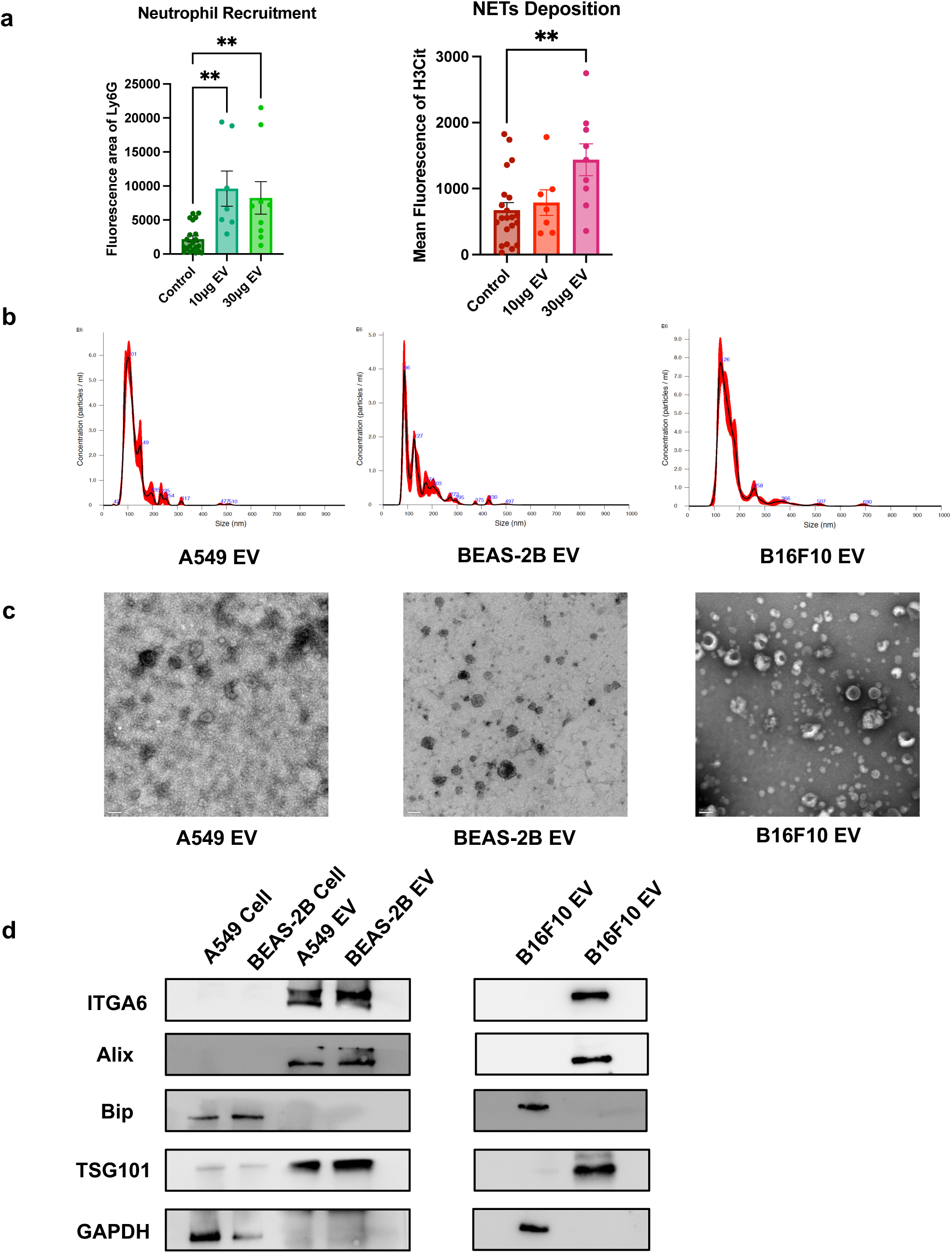
(a) Quantification of the area of lymphatic neutrophils (Ly6G), NETs (H3Cit) in draining LNs in non-tumour bearing mice after footpad injection of PBS or different dose of B16F10 EV, corresponding to the pictures in Fig. 7f. n = 5 Data shown as mean ± SEM. **, P < 0.01 by One-Way ANOVA. (b) Nanoparticle tracking assay (NTA) showing the size distribution of EVs. (c) Representative transmission electron microscopy (TEM) images of EVs. Scale bars represent 100nm. (d) Representative Western Blot images of EV and cell lysates.

**Figure 1:**
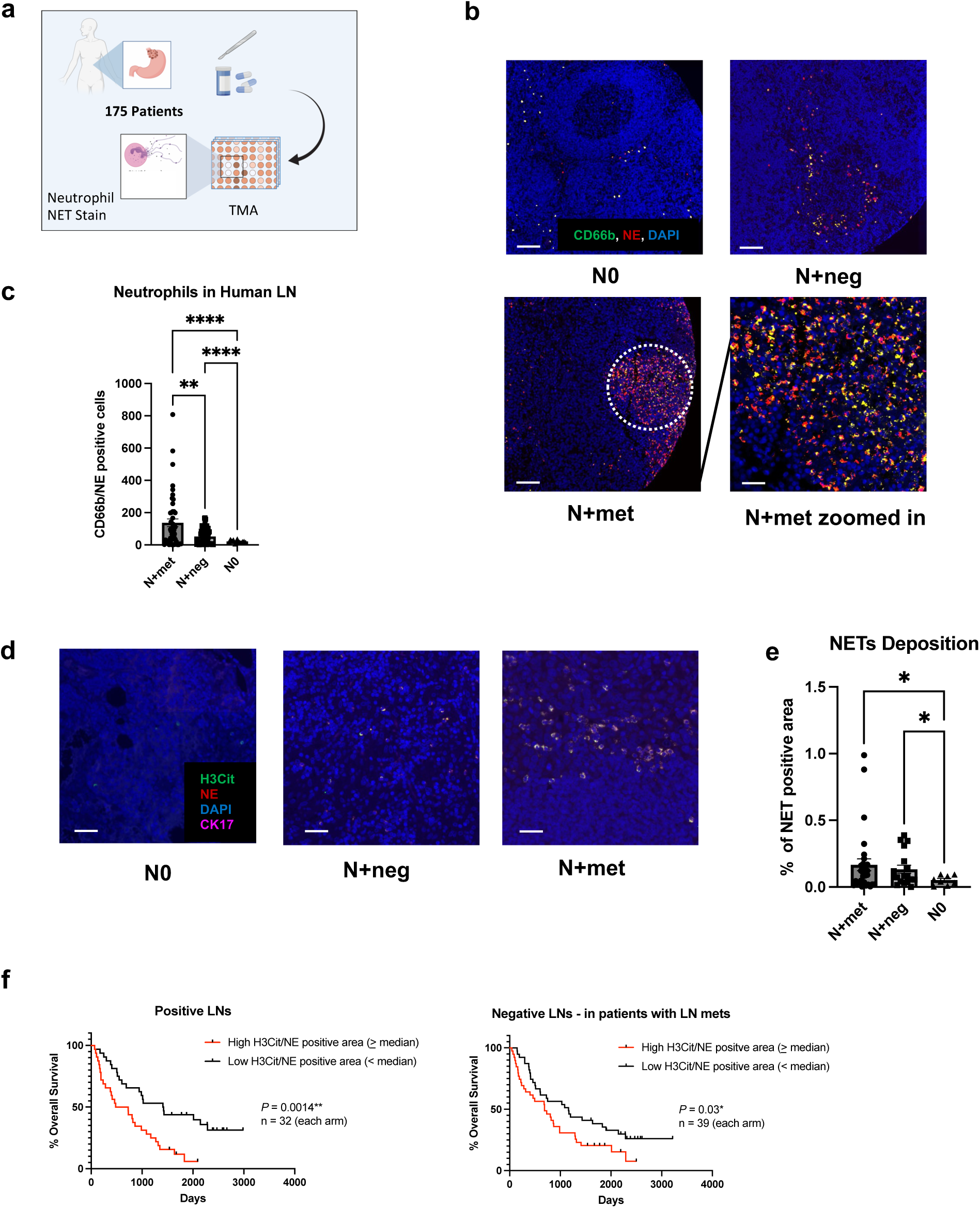
Neutrophils and NETs in regional lymph nodes (LNs) of gastroesophageal adenocarcinoma (GEA) patients. (a) Schematic illustration of the TMA constructions. Sample size: LNs from node negative patients (N0) =13, tumour negative LNs from node positive patients (N+neg) =91 and tumour positive LNs (N+met) =71. Each patient sample corresponds to one tissue core on TMA slides. (b) Representative images of neutrophils in different patient LNs. Scale bars represent 100μm. White circle indicates the zoomed area shown in bottom right. Scale bars represent 50μm. (c) Quantification of LN neutrophils (CD66b/NE double positive cells) per TMA core for N+met, N+neg and N0 LNs. (d) Representative images of NETs deposition pattern in N+met, N+neg and N0 LNs. Scale bars represent 50μm. (e) Quantification of %NETs positive area per core in N+met, N+neg and N0 LNs. Data shown as mean ± SEM. *, P < 0.05; **, P < 0.01; **** P < 0.001 by Brown-Forsythe ANOVA test. (f) Kaplan-Meier survival curves comparing survival of nodal positive GEA patients with low versus high levels of median lymphatic NETs positive area, in both tumour positive and negative LNs. P value by Log-rank (Mantel-Cox) test.

**Figure 2:**
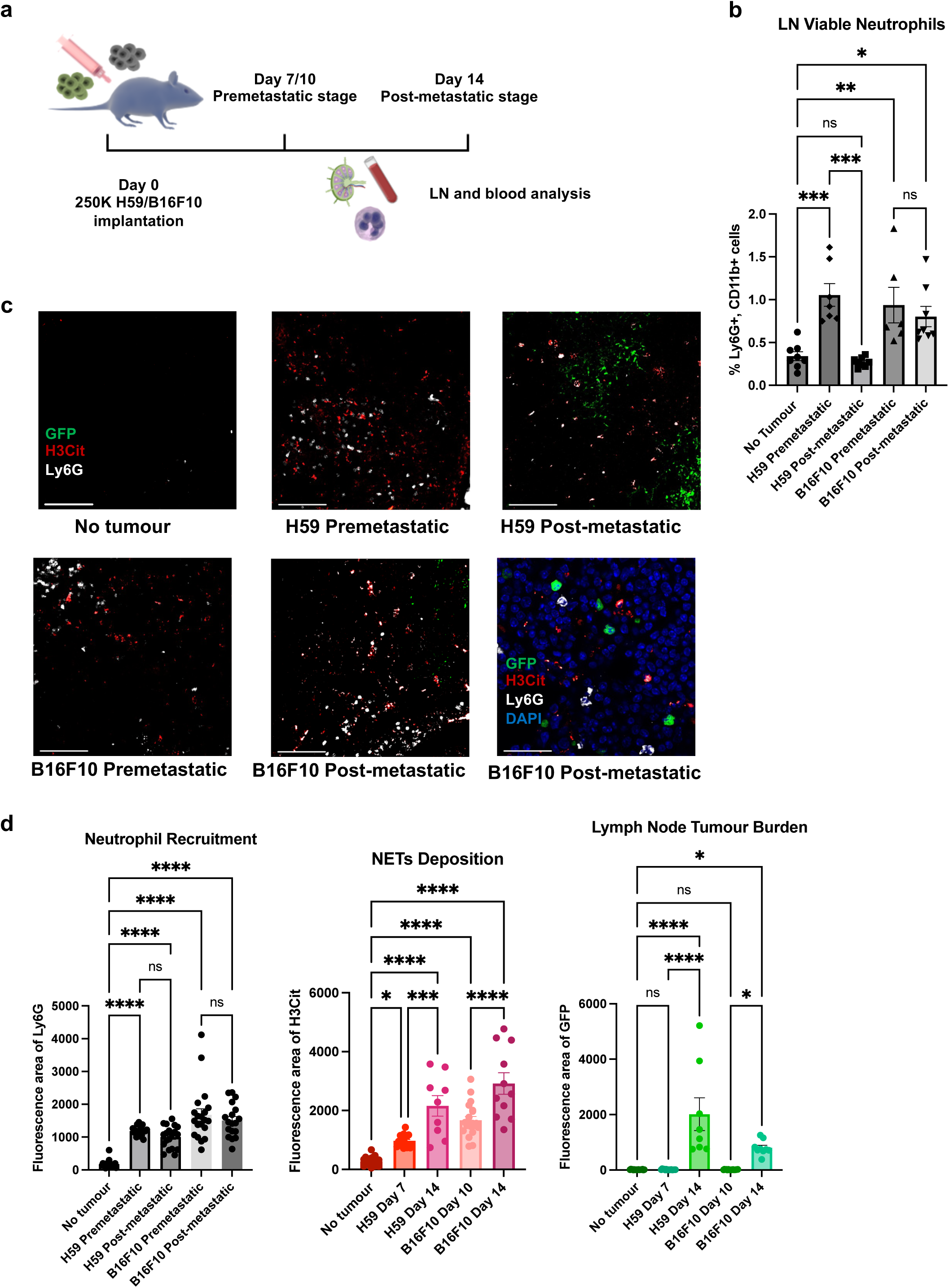
The dynamic of LN neutrophil recruitment and NET deposition in mouse model. (a) Schematic illustration of the animal model construction. 250k H59 or B16F10 cells were injected into the flank of C57bl/6 mice (b) Percentage of LN viable neutrophils within all viable LN cells on a time course by flow cytometry. (c) Representative images of LNs on a time course for both H59 lung cancer and B16F10 melanoma. Scale bars represent 100μm. Zoomed NETs details are shown in bottom right. Scale bars represent 20μm. (d) Quantification of the area of LN neutrophils (Ly6G), NETs (H3Cit) and tumour (GFP). Each data point is an image analysed, mouse n =10. Data shown as mean ± SEM. *, P < 0.05; **, P < 0.01; ***, P < 0.005; **** P < 0.001 by One-Way ANOVA.

**Figure 3:**
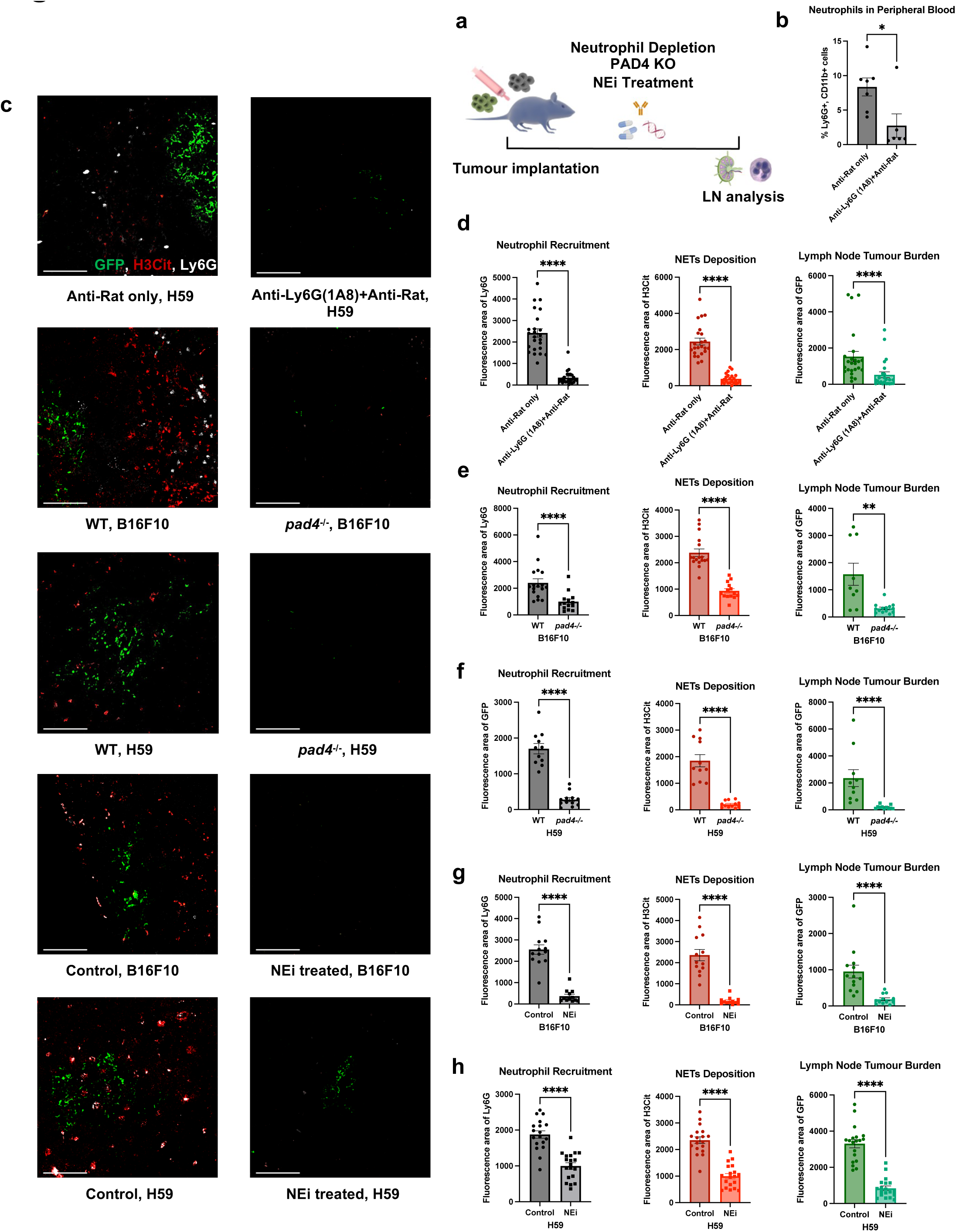
Neutrophil depletion and NETs inhibition abolishes LN metastasis. (a) Schematic illustration of the treatments in animal models. (b) Flow cytometry on blood leukocyte indicates sufficient depletion of neutrophils. (c) Representative images of tumour draining LNs on day 14 post tumour inoculation for both H59 and B16F10, comparing wildtype/no treatment mice to neutrophil-depleted, *pad4* knockout or NEi treated mice. Scale bars represent 100μm. (d)-(h) Quantification of the area of neutrophils (Ly6G), NETs (H3Cit) and metastasis (GFP) in fay 14 LNs for both H59 and B16F10, comparing wildtype/no treatment mice to neutrophil-depleted, *pad4* knockout or NEi treated mice. Each data point is an image analysed, mouse n =10. Data shown as mean ± SEM. *, P < 0.05; **, P < 0.01; ***, P < 0.005; **** P < 0.001 by Mann-Whitney t test.

**Figure 4:**
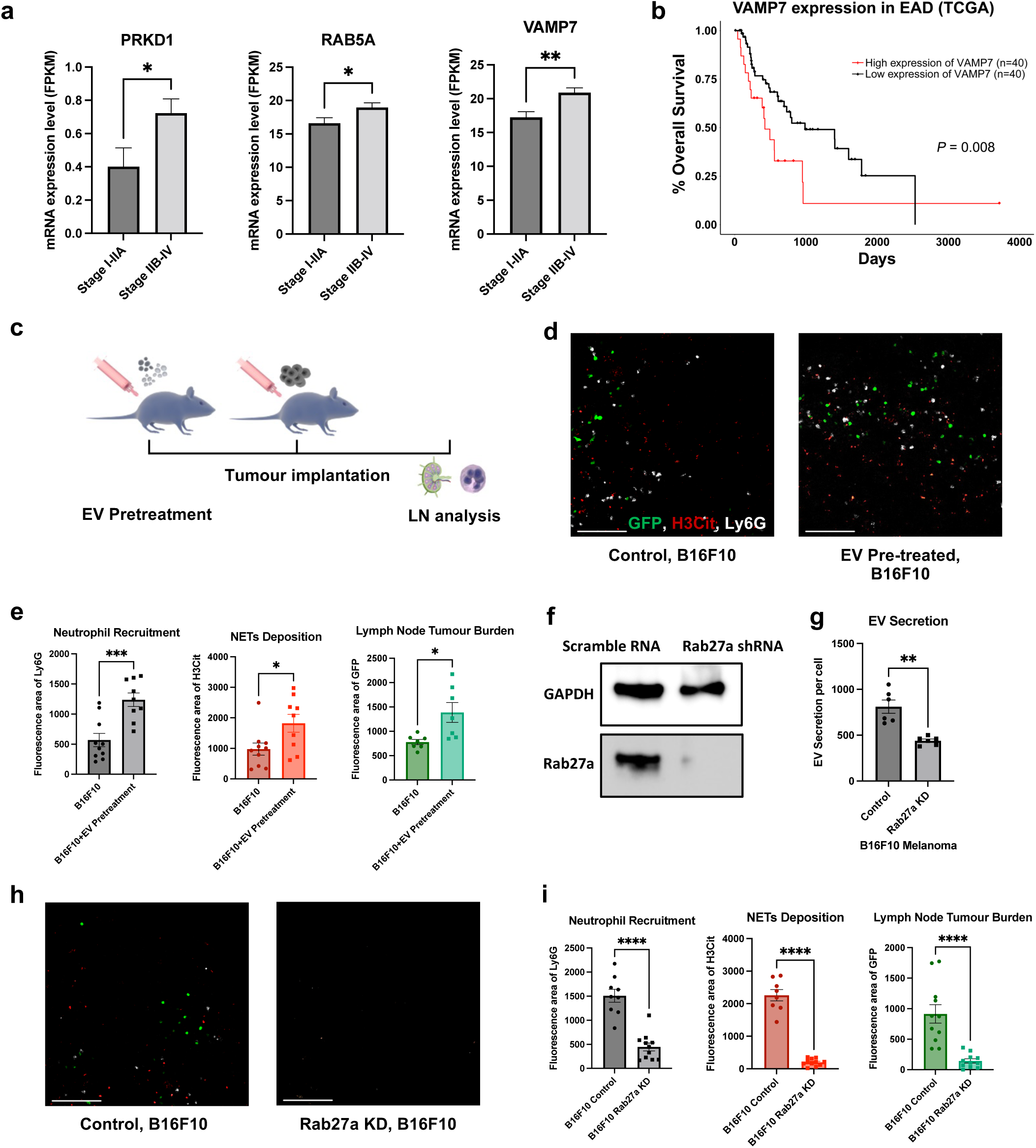
EVs are essential for LNs NETs formation and metastasis. (a) Statistical analysis of TCGA mRNA data of EV synthesis related genes in primary esophageal adenocarcinoma (EAD) tumour tissue. n = 15 (Stage I-IIA), n = 65 (Stage IIb-IV). (b) Kaplan-Meier survival curves comparing the survival of EAD patients with low versus high levels of VAMP7 mRNA in tumour. (c) Schematic illustration of EV pre-treatment in animal models. (d) Representative images of tumour draining LNs on day 14 post tumour inoculation for B16F10, with or without pre-treatment of EVs. Scale bars represent 100μm. (e) Quantification of the area of neutrophils (Ly6G), NETs (H3Cit) and tumour (GFP) in day 14 LN for B16F10, with or without EV pre-treatment. (f) Representative Western Blot images indicating the knockdown of *Rab27a* expression in B16F10 cells. (g) Nanoparticle Tracking Analysis (NTA) indicates *Rab27a* knockdown in B16F10 cells leads to decreased EV secretion. (h) Representative images of tumour draining LNs on day 14 post tumour inoculation for B16F10, comparing mice injected with B16F10 infected with lentivirus containing scramble RNA or *Rab27a* shRNA. Scale bars represent 100μm. (i) Quantification of the area of neutrophils (Ly6G), NETs (H3Cit) and metastasis (GFP) in day 14 LNs for B16F10, comparing mice injected with B16F10 infected with lentivirus contain scramble RNA or *Rab27a* shRNA. Each data point is an image analysed, mouse n =10. Data shown as mean ± SEM. *, P < 0.05; **, P < 0.01; ***, P < 0.005; **** P < 0.001 by Mann-Whitney t test.

**Figure 5:**
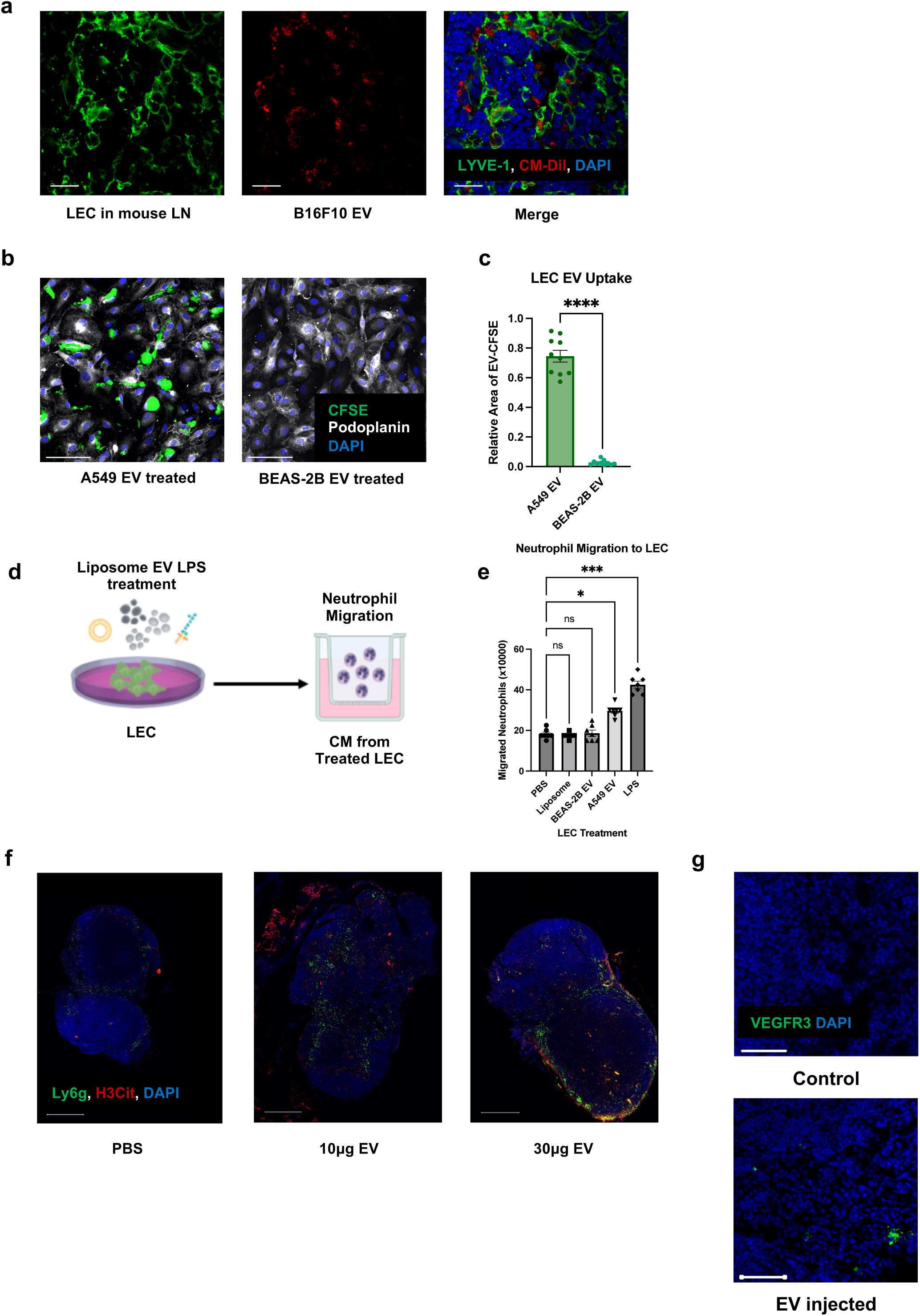
Lymphatic endothelial cells (LECs) are the LN recipient cells of EVs. LECs regulate neutrophil recruitment and NETs formation. (a) Representative images of draining LNs after footpad B16F10 EV injection, showing LECs uptaking EVs. Scale bars represent 20μm. (b) Representative images of LECs after A549 and BEAS-2B EV treatment. Scale bars represent 100μm. (c) Quantification of the EV uptake (relative fluorescence area of CFSE to DAPI) in LECs. (d) Schematic illustration of the Boyden chamber transwell assay. (e) Quantification of Boyden chamber transwell assay, showing neutrophil migration towards CM from treated LECs. Data shown as mean ± SEM. *, P < 0.05; ***, P < 0.005; **** P < 0.001 by Mann-Whitney t test or One-Way ANOVA. (f) Representative images of draining LNs after footpad PBS or different doses of B16F10 EV injection, indicating the subsequent neutrophil recruitment and NETs deposition. Scale bars represent 500μm. (g) Representative images of draining LNs after footpad PBS or 10μg B16F10 EV injection, indicating the increased expression of premetastatic marker VEGFR3. Scale bars represent 50μm

**Figure 6:**
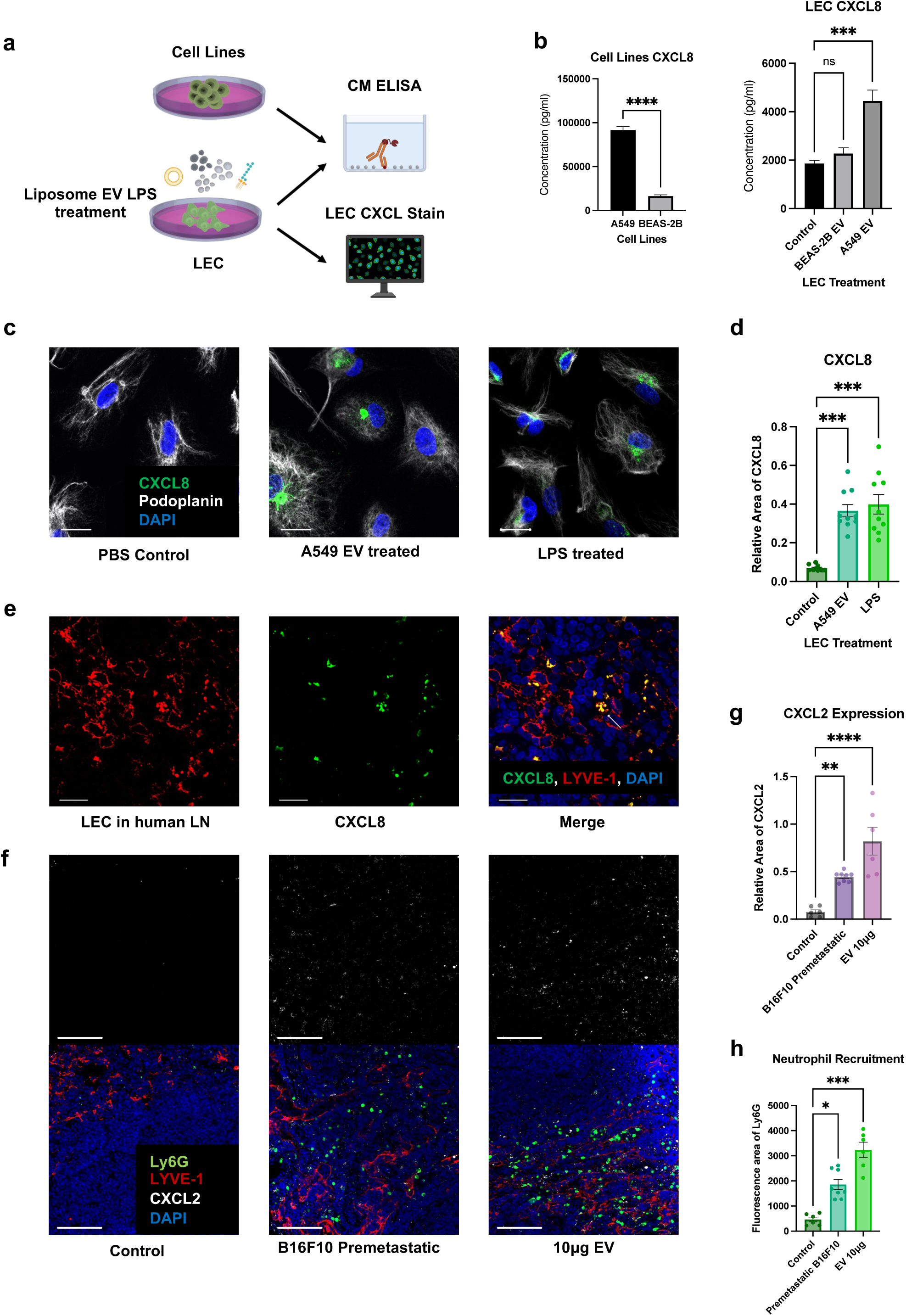
LECs secrete neutrophil chemoattractants and NETs inducers upon EV uptake. (a) Schematic illustration of the ELISA and LEC CXCL8 stain. (b) ELISA of CXCL8 level in conditioned media (CM) of A549, BEAS-2B and EV-treated LECs. (c) Representative images of LECs expressing CXCL8 after different kinds of treatments. Scale bars represent 20μm. (d) Quantification of CXCL8 expression in LECs (relative fluorescence area of CXCL8 to DAPI). (e) Representative images of LECs expressing CXCL8 in tumour positive nodes of lung cancer patients. Scale bars represent 20μm. (f) Representative images of LECs expressing CXCL2 and recruiting neutrophils mouse LNs. Scale bars represent 100μm. (g) and (h) Quantification of CXCL2 expression (relative fluorescence area of CXCL2 to LYVE-1) in LECs and neutrophil recruitment (Ly6G fluorescence area) in mouse LNs. Each data point is an image analysed, mouse n =10. Data shown as mean ± SEM. *, P < 0.05; **, P < 0.01; ***, P < 0.005; **** P < 0.001 by Mann-Whitney t test, One-Way ANOVA or Kruskal-Wallis test.

**Figure 7:**
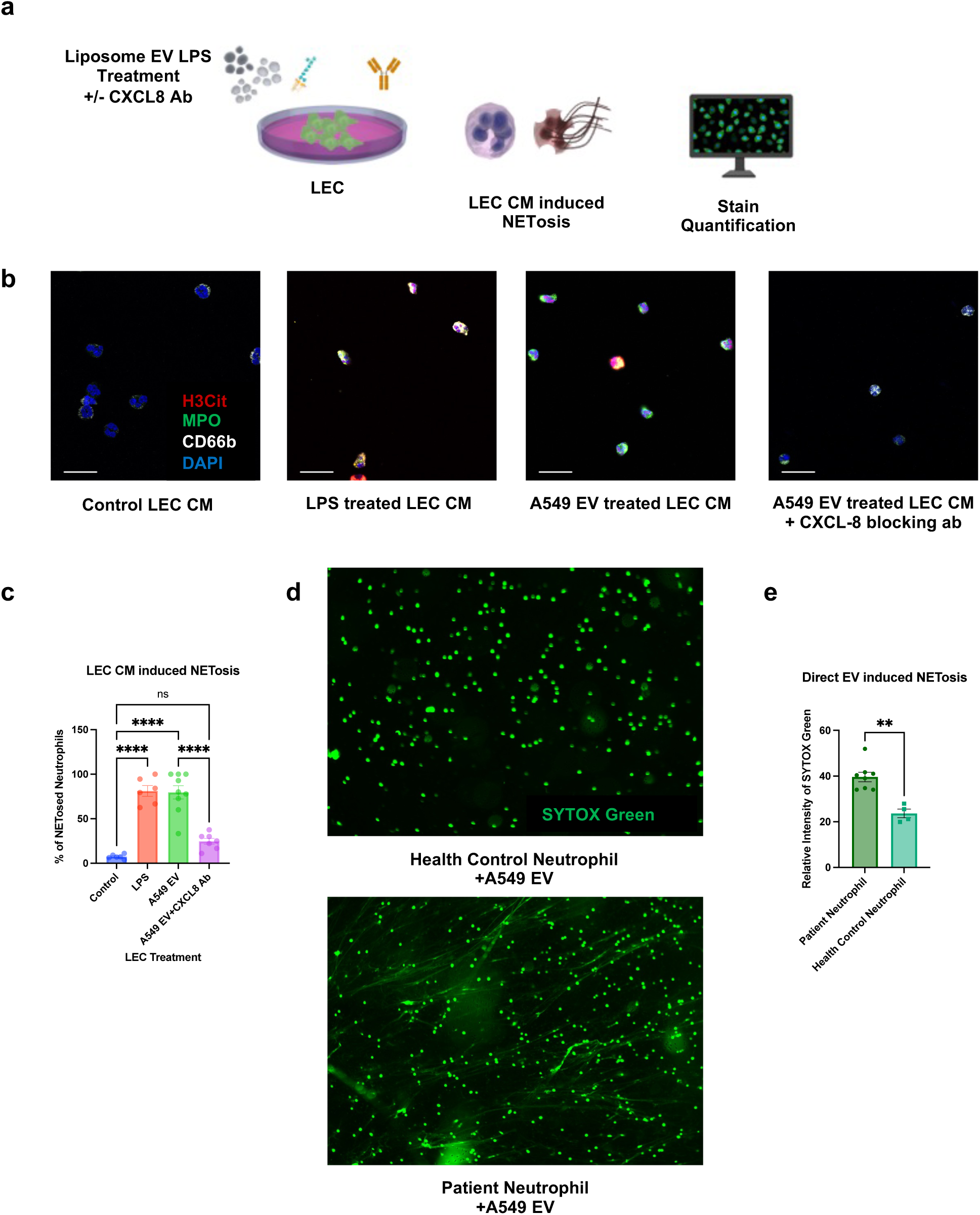
Lymphatic CXCL8 secretion upon EV uptake leads to NETs formation. (a)Schematic illustration of the *in vitro* NETosis assay induced by CM of treated LECs. (b) Representative images of neutrophils treated with CM from treated LECs. Scale bars represent 20μm. (c) Quantification of LEC CM induced NETosis (% of H3Cit positive neutrophils). (d) Representative images of indicating the different proneness to form NETs for neutrophils from health control and GEA patients after A549 EV treatment. Scale bars represent 100μm. (d) Quantification of propensity of neutrophil to form NETs induced by EVs (Relative fluorescence intensity of SYTOX Green treated to untreated neutrophil) from health control and GEA patients). Data shown as mean ± SEM.**, P < 0.01; ****, P < 0.001 by Kruskal-Wallis test or Mann-Whitney t test.

