## Supplementary figures and tables for "Tumour Extracellular Vesicles Induce Neutrophil Extracellular Traps To Promote Lymph Node Metastasis"

### Figure S1

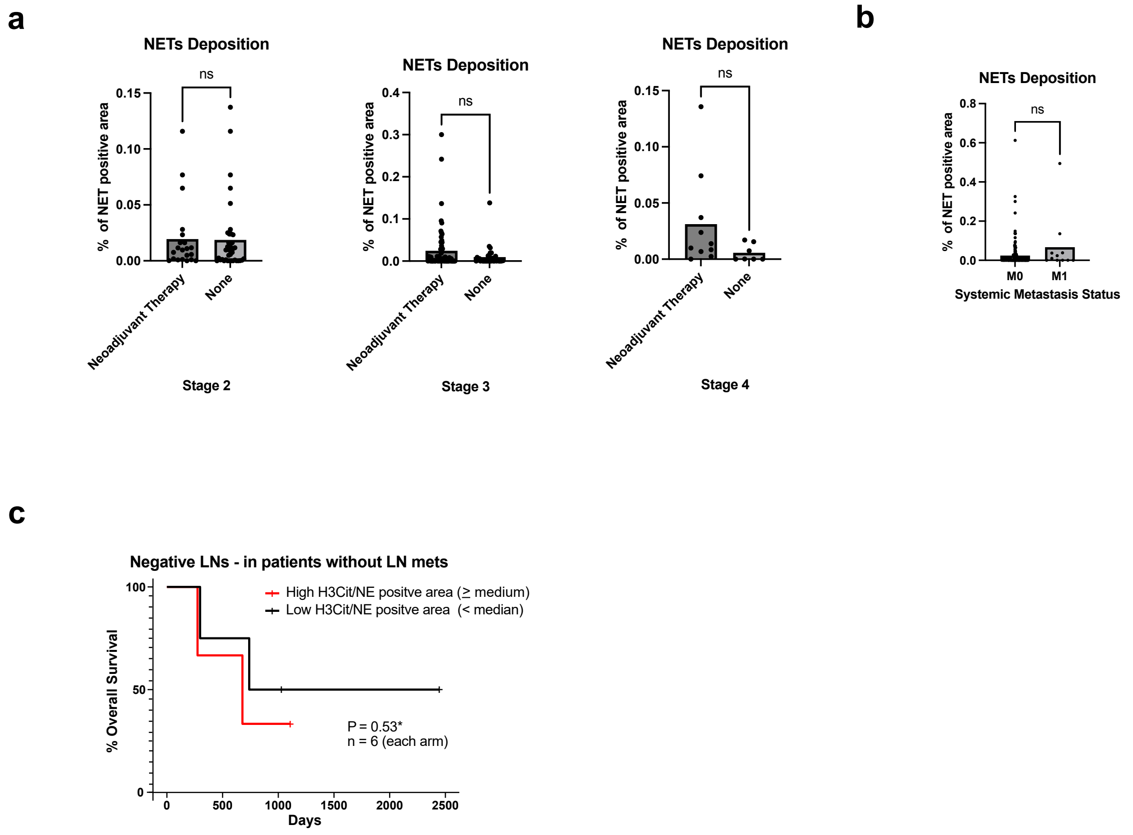
a
NETs Deposition
ns
% of NET positive area
0.15
0.10
0.05
0.00
Neoadjuvant Therapy
None
Stage 2
NETs Deposition
ns
% of NET positive area
0.4
0.3
0.2
0.1
0.0
Neoadjuvant Therapy
None
Stage 3
NETs Deposition
ns
% of NET positive area
0.15
0.10
0.05
0.00
Neoadjuvant Therapy
None
Stage 4
b
NETs Deposition
ns
% of NET positive area
0.8
0.6
0.4
0.2
0.0
M0
M1
Systemic Metastasis Status
c
Negative LNs - in patients without LN mets
High H3Cit/NE positve area (≥ medium)
Low H3Cit/NE positve area (< median)
% Overall Survival
100
50
0
P = 0.53*
n = 6 (each arm)
0
500
1000
1500
2000
2500
Days

### Figure S2

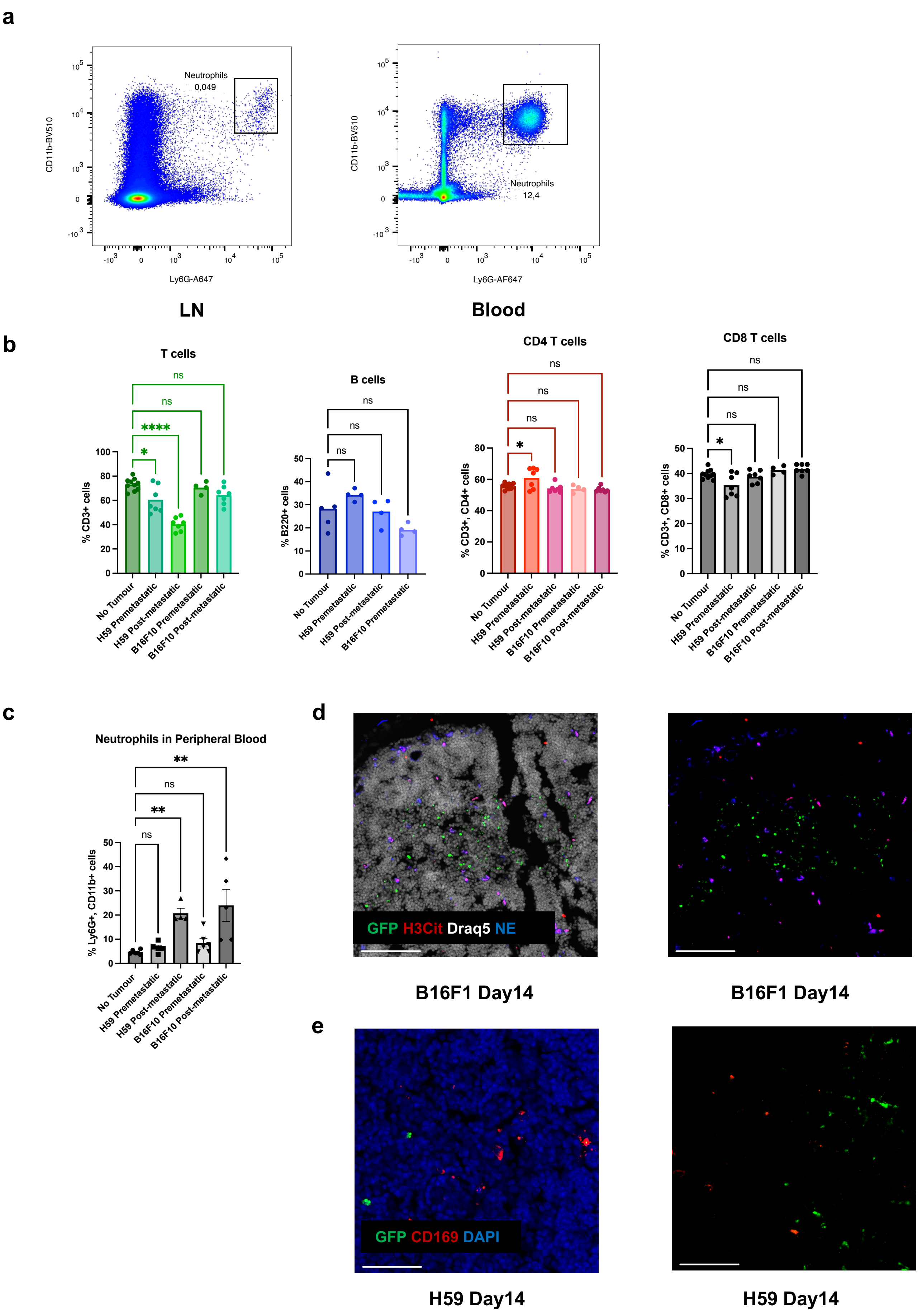

a
Neutrophils
0,049
CD11b-BV510
Ly6G-A647
LN
Neutrophils
12,4
CD11b-BV510
Ly6G-AF647
Blood
b
T cells
% CD3+ cells
B cells
% B220+ cells
CD4 T cells
% CD3+, CD4+ cells
CD8 T cells
% CD3+, CD8+ cells
ns
****
*
No Tumour
H59 Premetastatic
H59 Post-metastatic
B16F10 Premetastatic
B16F10 Post-metastatic
c
Neutrophils in Peripheral Blood
% Ly6G+, CD11b+ cells
**
ns
d
GFP H3Cit Draq5 NE
B16F1 Day14
B16F1 Day14
e
GFP CD169 DAPI
H59 Day14
H59 Day14

### Figure S3

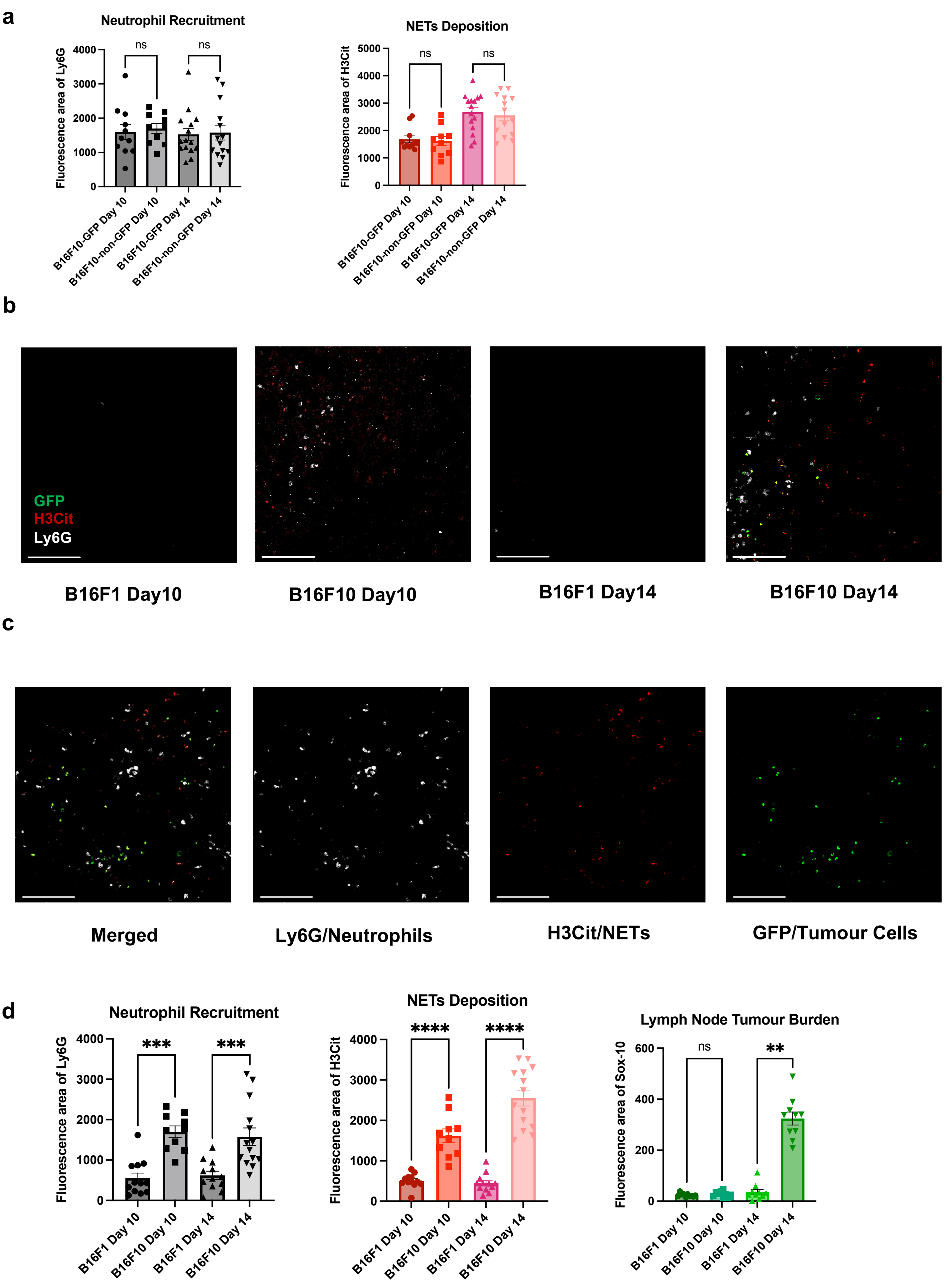

a
Neutrophil Recruitment
Fluorescence area of Ly6G
4000
3000
2000
1000
0
ns
ns
B16F10-GFP Day 10
B16F10-non-GFP Day 10
B16F10-GFP Day 14
B16F10-non-GFP Day 14
NETs Deposition
Fluorescence area of H3Cit
5000
4000
3000
2000
1000
0
ns
ns
B16F10-GFP Day 10
B16F10-non-GFP Day 10
B16F10-GFP Day 14
B16F10-non-GFP Day 14
b
GFP
H3Cit
Ly6G
B16F1 Day10
B16F10 Day10
B16F1 Day14
B16F10 Day14
c
Merged
Ly6G/Neutrophils
H3Cit/NETs
GFP/Tumour Cells
d
Neutrophil Recruitment
Fluorescence area of Ly6G
4000
3000
2000
1000
0
***
***
B16F1 Day 10
B16F10 Day 10
B16F1 Day 14
B16F10 Day 14
NETs Deposition
Fluorescence area of H3Cit
4000
3000
2000
1000
0
****
****
B16F1 Day 10
B16F10 Day 10
B16F1 Day 14
B16F10 Day 14
Lymph Node Tumour Burden
Fluorescence area of Sox-10
600
400
200
0
ns
**
B16F1 Day 10
B16F10 Day 10
B16F1 Day 14
B16F10 Day 14

### Figure S4

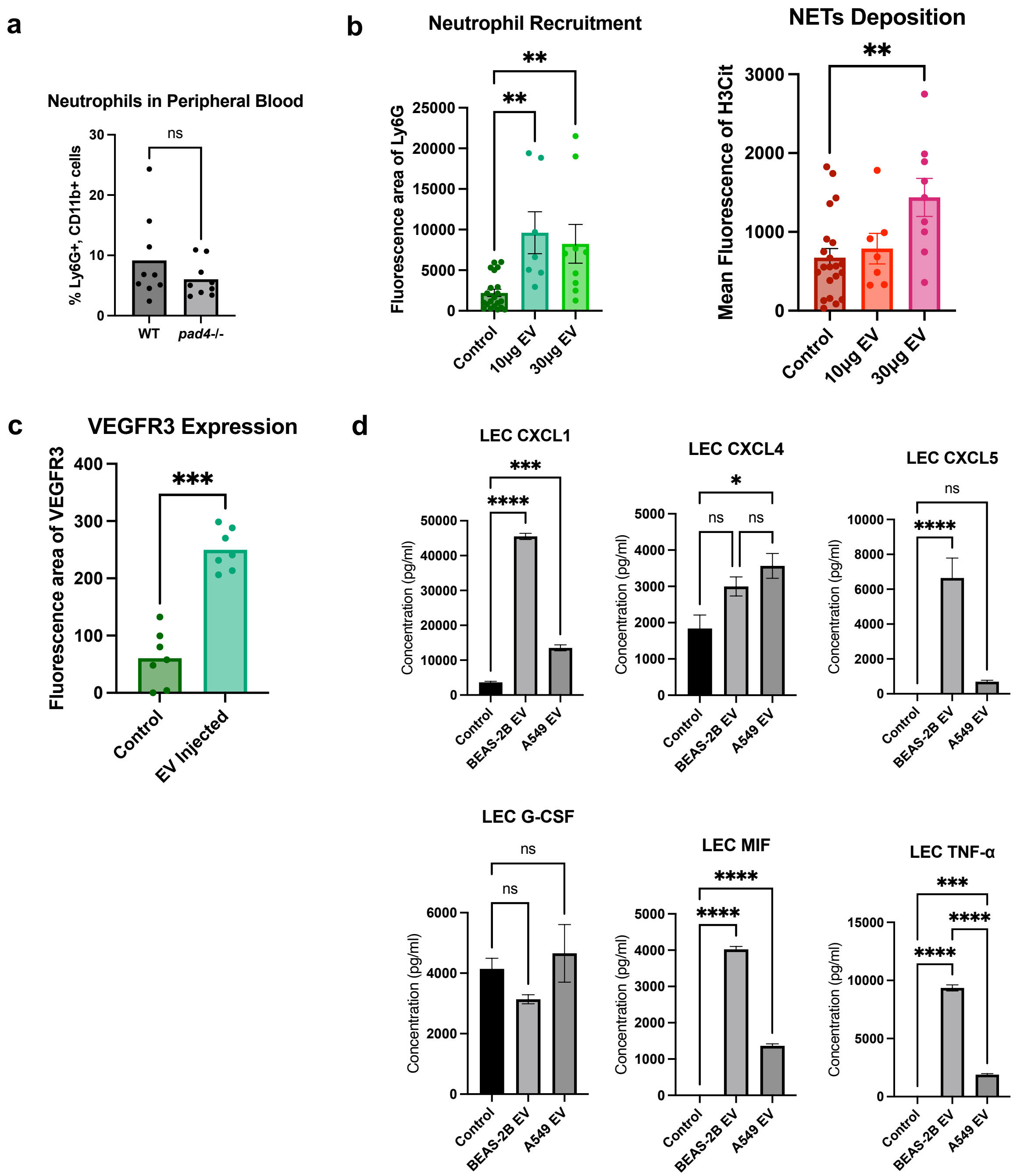
a
Neutrophils in Peripheral Blood
% Ly6G+, CD11b+ cells
ns
WT
pad4-/-
b
Neutrophil Recruitment
Fluorescence area of Ly6G
**
Control
10μg EV
30μg EV
NETs Deposition
Mean Fluorescence of H3Cit
c
VEGFR3 Expression
Fluorescence area of VEGFR3
***
Control
EV Injected
d
LEC CXCL1
LEC CXCL4
LEC CXCL5
LEC G-CSF
LEC MIF
LEC TNF-α
Concentration (pg/ml)
BEAS-2B EV
A549 EV
****
*

### Figure S5

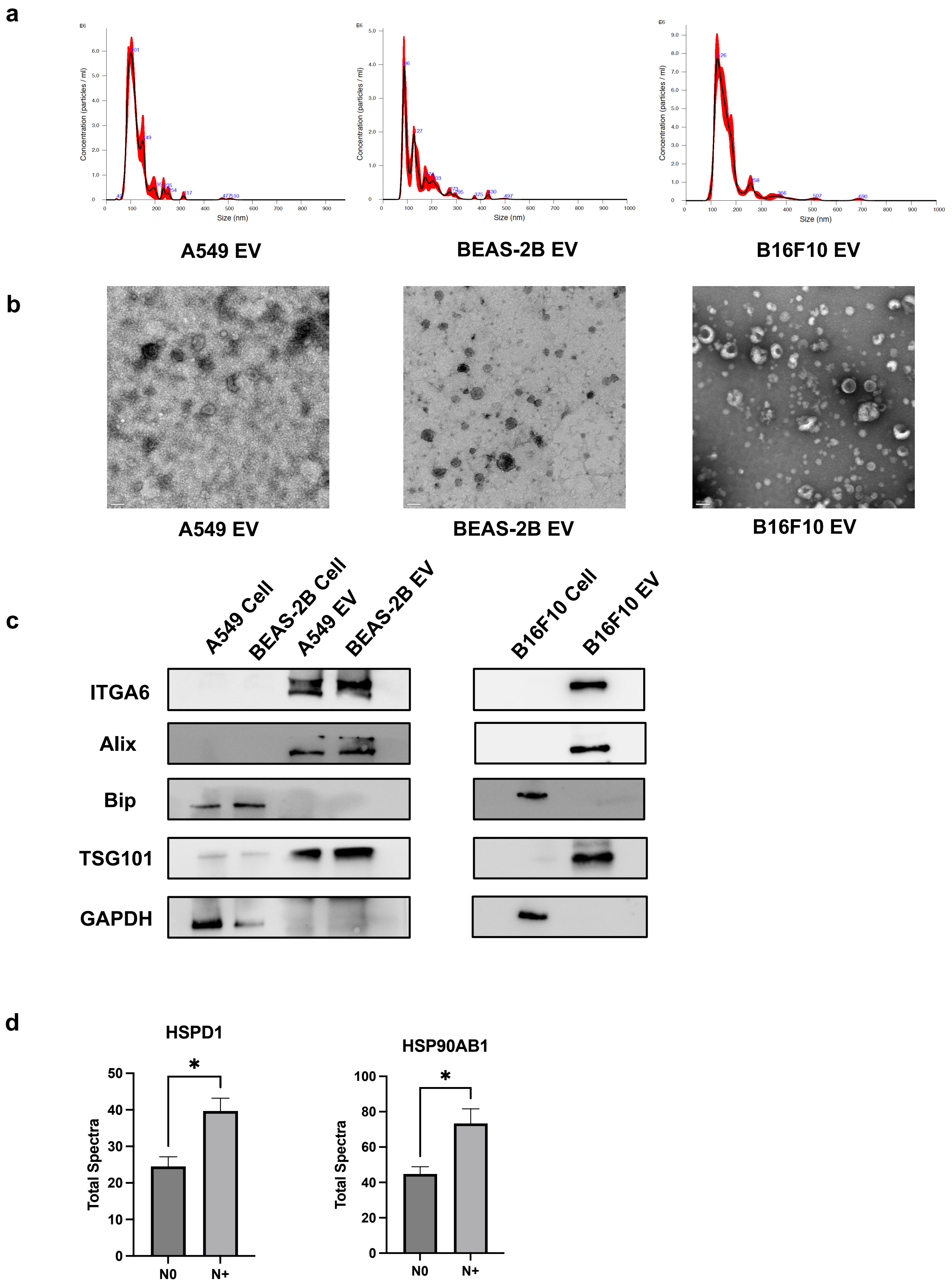
a
Concentration (particles / ml)
Size (nm)
A549 EV
BEAS-2B EV
B16F10 EV
b
A549 EV
BEAS-2B EV
B16F10 EV
c
A549 Cell
BEAS-2B Cell
A549 EV
BEAS-2B EV
B16F10 Cell
B16F10 EV
ITGA6
Alix
Bip
TSG101
GAPDH
d
HSPD1
HSP90AB1
Total Spectra
*
N0
N+

**Figure S1:** (a) Quantification of %NETs positive area per LN core, divided by stage and comparing patients who received neoadjuvant therapies or not. (Stage 2: n=21 vs. 38, Stage 3: n=64 vs. 33, Stage 4: n=10 vs 7) (b) Quantification of %NETs positive area per LN core in M0 (n=159) and M1 (n=11) patients. Data shown as mean ± SEM. NS, $P > 0.05$ by Mann-Whitney t test. (c) Kaplan-Meier survival curves comparing survival of N0 GEA patients with low versus high levels of median lymphatic NETs positive area. P value by Log-rank (Mantel-Cox) test.

**Figure S2:** (a) Flow cytometry gating strategies of mouse LN and blood neutrophils. (b) Percentage of mouse LN T cells, B Cells, CD4+ T cells and CD8+ T cells within all LN cells on a time course by flow cytometry. (c) Percentage of neutrophils within blood leukocytes on a time course by flow cytometry. ns, $P > 0.05$; *, $P < 0.05$; **, $P < 0.01$; **** $P < 0.001$ by One-Way ANOVA. (d) Representative images of post-metastatic LNs for B16F10 melanoma using a different staining panel. Scale bars represent 100µm. (e) Representative images of post-metastatic LNs for both H59 lung cancer, demonstrating that GFP signals do not colocalize with CD169+ macrophages. Scale bars represent 50µm.

**Figure S3:** (a) Quantification of the area of lymphatic neutrophils (Ly6G) and NETs (H3Cit) in pre- and post- metastatic LNs for both B16F10-non-GFP and B16F10-GFP cells. (b) Representative images of tumour draining LNs on a time course for both B16F1 and B16F10 melanoma cells. Scale bars represent 100µm. (c) Representative images of post-metastatic LNs for B16F10 melanoma with separated fluorescence panels. Scale bars represent 100µm. (d) Quantification of the area of lymphatic neutrophils (Ly6G), NETs (H3Cit) and tumour (Sox-10) in no tumour, pre- and post- metastatic nodes for both B16F1 and B16F10. Each data point is an image analysed; data shown as mean ± SEM. Mouse n = 10; **, $P < 0.01$; ***, $P < 0.005$; **** $P < 0.001$ by One-Way ANOVA.

**Figure S4:** (a) Baseline percentage of neutrophils within blood leukocytes for wildtype and PAD4 knockout mice. ns, $P > 0.05$ by t-test. (b) Quantification of neutrophils and NETs in mouse draining LNs after footpad PBS or different doses of B16F10 EV injection. (c) Quantification of VEGFR3 in mouse draining LNs after footpad PBS or 10μg B16F10 EV injection. Each data point is an image analysed; mouse n = 5. (d) ELISA of CXCL1/4/5, G-CSF, MIF and TNF-α levels in conditioned media (CM) of LECs treated with A549 or BEAS-2B EV. Data shown as mean ± SEM; ns, $P > 0.05$; *, $P < 0.05$; **, $P < 0.01$; ***, $P < 0.005$; **** $P < 0.001$ by Mann-Whitney t test or One-Way ANOVA.

**Figure S5:** (a) Nanoparticle tracking assay (NTA) showing the size distribution of EVs. (b) Representative transmission electron microscopy (TEM) images of EVs. Scale bars represent 100nm. (c) Representative Western Blot images of EV and cell lysates. (d) Quantification of total protein spectra count of HSPD1 and HSP90AB1 in plasma EVs from GEA patients with or without nodal metastasis. n=4 for N0 patients and n=6 for N+ patients. *, $P < 0.05$ by Mann-Whitney t test.

Table S1 Demographic and clinical characteristics of patients with gastroesophageal adenocarcinoma in the TMA study

| **Patients with gastroesophageal adenocarcinoma in the TMA study (n = 175)** | |
|---|---|
| Mean Age, years | 68.08 |
| Sex, male | 78.28% |
| Neoadjuvant Treatment, yes | 54.48% |
| Clinical Stage 1 | 21.65% |
| Clinical Stage 2 | 31.85% |
| Clinical Stage 3 | 36.94% |
| Clinical Stage 4 | 9.55% |
| Clinical T1 | 12.10% |
| Clinical T2 | 21.65% |
| Clinical T3 & 4 | 66.24% |
| Clinical N, yes | 55.69% |
| Clinical M, yes | 10.82% |
| Pathologic Stage 1 | 18.07% |
| Pathologic Stage 2 | 22.29% |
| Pathologic Stage 3 | 51.20% |
| Pathologic Stage 4 | 8.43% |
| Pathologic T1 | 14.94% |
| Pathologic T2 | 19.55% |
| Pathologic T3 & T4 | 65.51% |
| Pathologic N0 | 25.28% |
| Pathologic N1 | 19.54% |
| Pathologic N2 | 26.43% |
| Pathologic N3 | 28.75% |
| Pathologic M, yes | 8.82% |
| Recurrence, yes | 39.65% |
| Grade, poor | 52.35% |
| Grade, moderate | 41.76% |
| Grade, well | 5.88% |
| Lymphovascular Invasion, yes | 70.00% |
| Median Overall Survival (days) | 678 |
| Median Disease-Free Survival (days) | 1064 |

Table S2 Demographic and clinical characteristics of patients with oesophageal adenocarcinoma from TCGA

| **Patients with esophageal adenocarcinoma from TCGA (n =80)** | |
|---|---|
| Mean Age, years | 66.2 |
| Sex, male | 13.75% |
| Clinical Stage 1 | 2.50% |
| Clinical Stage 2 | 3.75% |
| Clinical Stage 3 | 7.50% |
| Clinical Stage 4 | 6.25% |
| Clinical Stage, NA | 80.00% |
| Clinical T1 | 1.25% |
| Clinical T2 | 2.50% |
| Clinical T3 | 13.75% |
| Clinical T, NA | 82.50% |
| Clinical N, yes | 12.50% |
| Clinical M, yes | 6.25% |
| Pathologic Stage 1 | 12.50% |
| Pathologic Stage 2 | 27.50% |
| Pathologic Stage 3 | 33.75% |
| Pathologic Stage 4 | 6.25% |
| Pathologic Stage, NA | 20.00% |
| Pathologic Stage, I_IIA | 18.75% |
| Pathologic Stage, IIB_IV | 61.25% |
| Pathologic T0 | 1.25% |
| Pathologic T1 | 26.25% |
| Pathologic T2 | 13.25% |
| Pathologic T3 & T4 | 43.75% |
| Pathologic T, NA | 15.00% |
| Pathologic N0 | 25.00% |
| Pathologic N1 | 46.25% |
| Pathologic N2 | 6.25% |
| Pathologic N3 | 6.25% |
| Pathologic N, NA | 16.25% |
| Pathologic M, yes | 6.25% |
| Median Overall Survival (days) | 393 |
| Status, dead | 41.25% |

Table S3 Demographic and clinical characteristics of patients with oesophageal adenocarcinoma for plasma EV proteomics study

| **Patients with esophageal adenocarcinoma from TCGA (n =10)** | |
|---|---|
| Mean Age, years | 67.5 |
| Sex, male | 90% |
| Clinical Stage 1 | 0% |
| Clinical Stage 2 | 0% |
| Clinical Stage 3 | 90% |
| Clinical Stage 4 | 10% |
| Clinical T1 | 10% |
| Clinical T2 | 0% |
| Clinical T3 | 90% |
| Clinical N, yes | 40% |
| Clinical M, yes | 20% |

Table S4 Resources table of antibodies, reagents, cells and viruses.

| REAGENT or RESOURCE | SOURCE | IDENTIFIER |
|---|---|---|
| Antibodies | | |
| anti-human-NE | R&D Systems | MAB91671100 |
| anti-human-CD66b | BioLegend | 392902 |
| anti-mouse/human-H3Cit | abcam | ab5103 |
| anti-human-Cytokeratin 7 | abcam | ab9021 |
| anti-Mouse-Ly6G (1A8) | BioXCell | BE0075-1 |
| anti-Rat-kappa light chain (MAR18.5) | BioXCell | BE0122 |
| Mouse Fc-block | BD | 553141 |
| anti-mouse-CD45.2- eFL450 | Invitrogen | 48-0454-80 |
| anti-mouse-CD11b-BV510 | BioLegend | 101263 |
| anti-mouse-Ly6G-AF647 | Biolegend | 127610 |
| anti-mouse-SOX-10-AF488 | Novus Bio | NBP2-59621AF488 |
| anti-mouse-LYVE-1 | R&D Systems | AF2125 |
| anti-mouse/human-NE | Bioss | bs-6982R |
| anti-mouse-Ly6G-FITC | Biolegend | 11-9668-80 |
| anti-mouse-MIP2/CXCL2 | PeproTech | 500-P130 |
| goat-anti-rabbit-AF568 | Invitrogen | A11011 |
| donkey-anti-goat-AF488 | Invitrogen | A110555 |
| donkey-anti-goat-AF568 | Invitrogen | A11057 |
| donkey-anti-rabbit-AF647 | Invitrogen | A-31573 |
| anti-human-LYVE-1 | R&D System | AF2089 |
| anti-human-CXCL8 | R&D System | MAB208 |
| donkey-anti-mouse-AF488 | Invitrogen | A-21202 |

| | | |
|---|---|---|
| anti-mouse/human-ITGA6 | Cell Signalling | 3750 |
| anti-mouse/human-Alix | Cell Signalling | 2171 |
| anti-mouse/human BiP | Cell Signalling | 3183 |
| anti-mouse-Rab27a | Cell Signalling | 69295 |
| anti-mouse/human-GAPDH | Invitrogen | MA5-15738 |
| anti-human-Podoplanin-AF647 | Biolegend | 916609 |
| anti-human-CXCL8-AF488 | R&D System | IC208G |
| anti-human-MPO-FITC | abcam | ab11729 |
| anti-human-CD66b-APC | Miltenyl Biotech | 130-122-966 |
| goat-anti-rabbit-AF568 | Invitrogen | A11011 |
| Virus strains | | |
| Murine lentivirus with Rab27a-mouse shRNA Sequence: CTGGATAAGCCAGCTACAGATGCACGCGT | Origene | MR202531L3V |
| Murine lentivirus with control scramble RNA | Origene | PS100092V |
| Chemicals, peptides, and recombinant proteins | | |
| Opal 6-Plex Manual Detection Kit | Akoya Biosciences | NEL811001KT |
| DAPI | Invitrogen | D21490 |
| Neutrophil elastase inhibitor Sivelestat | abcam | ab146184 |
| Viability Dye Eflour780 | eBioscience | 65-0865-14 |
| DRAQ5 Fluorescent Probe Solution | Thermo Scientific | 62251 |
| CellTrace CFSE Cell Proliferation Kit | Invitrogen | C34554 |
| CellTracker CM-DiI Dye | Invitrogen | C7000 |
| Laemmli SDS sample buffer | BIO-RAD | 1610747 |
| SYTOX Green ( | Invitrogen | |
| 100nm Liposome control | Encapsula Nanoscience | CEP-500 |
| Critical commercial assays | | |
| Transwell Permeable Supports | Costar | 3421 |
| Multiplex ELISA kit | RayBioTech | QAH-CUST-H10 |
| Experimental models: Cell lines | | |
| B16-F10 | Dr. John Stagg Lab, originally purchased from ATCC | CRL-6475 |
| B16-F1 | Dr. Ian Watson Lab, originally purchased from ATCC | CRL-6323 |
| H59 | Generated by Dr. Pnina Brodt Lab | / |
| A549 | ATCC | CCL-185 |
| BEAS-2B | ATCC | CRL-9609 |
| LEC | PromoCell | C-12216 |
| Experimental models: Organisms/strains | | |

| | | |
|---|---|---|
| Mouse: C57BL/6NCrl | Charles River Laboratories | C57BL/6NCrl |
| Mouse: PAD4 knockout (pad4–/–, C57BL/6NCrl background) | Dr. Alan Tsung Lab | |
| Software and algorithms | | |
| FlowJo software | Tree Star | / |
| HALO | Indica Labs | / |
| ImageJ Software | NIH | / |
| GraphPad | Prism | / |
| RStudio (Version 3.6.3) | Posit.co | |
| Other | | |
| BD Pharm Lyse lysing solution | BD Pharm | 555899 |
| Mini-PROTEAN TGX Gels | BIO-RAD | 4561081 |
| Nitrocellulose membrane | BIO-RAD | 17094159 |
| Lymphocyte Separation Media | Wisent Bioproducts | 305-010-CL |
